# JunB controls intestinal effector programs in regulatory T cells

**DOI:** 10.1101/772194

**Authors:** Joshua D. Wheaton, Maria Ciofani

## Abstract

Foxp3-expressing regulatory T (Treg) cells are critical mediators of immunological tolerance to both self and microbial antigens. Tregs activate context-dependent transcriptional programs to adapt effector function to specific tissues; however, the factors controlling tissue-specific gene expression in Tregs remain unclear. Here, we find that the AP-1 transcription factor JunB regulates the intestinal adaptation of Tregs by controlling select gene expression programs in multiple Treg subsets. Treg-specific ablation of JunB results in immune dysregulation characterized by enhanced colonic T helper cell accumulation and cytokine production. However, in contrast to its classical binding-partner BATF, JunB is dispensable for maintenance of effector Tregs as well as most specialized Treg subsets. In the Peyer’s patches, JunB activates a transcriptional program facilitating the maintenance of CD25^-^ Tregs, leading to the complete loss of T follicular regulatory cells in the absence of JunB. This defect is compounded by loss of a separate effector program found in both major colonic Treg subsets that includes the cytolytic effector molecule granzyme B. Therefore, JunB is an essential regulator of intestinal Treg effector function through pleiotropic effects on gene expression.

## Introduction

The maintenance of immunological homeostasis is largely dependent on Foxp3^+^ regulatory T (Treg) cells, which suppress antigen-specific immune responses against both self and commensal microbial antigens. While Tregs control immune responses in part by inhibiting T cell priming in the spleen and lymph nodes (LN), it is now accepted that Tregs play wide-ranging roles in maintenance of organismal homeostasis through a myriad of activities in non-lymphoid tissues. These include regulation of insulin sensitivity in the visceral adipose tissue (VAT); control of wound-healing and repair responses in the skin, lung, and intestine; modulation of antibody responses in germinal centers (GCs); and suppression of adaptive immune responses against both dietary and microbial antigens in the intestine (1–3). Elucidating the factors affecting tissue-specific Treg function are thus critical to understanding both normal and adverse physiology of diverse organ systems.

Because of the range of conditions in which they operate, Tregs must adopt organ-specific patterns of gene expression that confer unique functional programs. Upon initial antigen recognition in the spleen and LN, resting central Treg (cTreg) cells upregulate an activation-dependent gene signature leading to acquisition of the effector Treg (eTreg) cell phenotype, which is characterized in part by loss of the lymphoid-homing receptors CD62L and CCR7 (1). Activated eTregs are then competent to undergo further organ-specific changes in gene expression – likely resulting from signals encountered both in the LN at the time of activation, and also after arrival in the target non-lymphoid tissue (4–6) – ultimately leading to unique tissue-specific transcriptional programs. However, despite substantial advances in our understanding of the diverse tissue-specific functions of Tregs, the transcriptional regulatory circuitry responsible for this diversity remains poorly understood.

A major mechanism driving tissue-specific gene expression in Tregs is the controlled expression of specific transcription factors (TFs). These include not only Treg subset-specific regulators – such as ROR*γ*t, Bcl-6, GATA-3, or PPAR*γ* – but also a large number of TFs with undefined roles in Treg biology (4, 6, 7). Previous studies have shown that expression of many TFs from the basic leucine zipper (bZIP)/AP-1 family are enriched in Tregs from diverse non-lymphoid tissues (4, 6, 8); however, our understanding of the role of AP-1 in tissue-specific Treg function remains limited. BATF, the most well-studied AP-1 TF in T cells, was shown to be critical for both differentiation of VAT-resident Tregs (9) and intestinal migration of Tregs (10). However, these observations can be largely explained by a general requirement for BATF in the initial differentiation of eTregs, such that BATF expression is important for all tissue Tregs but does not convey tissue specificity of gene expression (11). Conversely, we and others recently showed that the bZIP TF c-Maf is essential for the differentiation of both follicular regulatory T (Tfr) cells and ROR*γ*t^+^ Tregs, but does not play a role in differentiation of either eTregs or VAT Tregs, demonstrating that bZIP TFs can play tissue or subset-restricted roles in Treg biology (12, 13). Nevertheless, outside of these limited examples, the function of the vast majority of bZIP/AP-1 TFs in Treg biology remains unexplored.

A recent analysis comparing the regulomes of tissue-resident Tregs found that bZIP/AP-1 binding sites were strongly enriched within genomic regions that gained accessibility in tissue-specific Tregs relative to those from lymphoid organs. Based on gene expression and motif analyses, it was proposed that several AP-1 TFs – including JunB and JunD – might be broadly important in regulating tissue-specific transcription in Tregs, particularly in the VAT and colon (6). Previously, we and others found that the AP-1 TF JunB was essential for the differentiation of inflammatory T-helper 17 (Th17) cells (14–16), demonstrating that JunB plays subset-restricted roles in T cell programming. More recently, JunB was reported to control differentiation and suppressive functions of eTregs (17); however, many of the conclusions from this study are confounded by the use of a CD4-cre-mediated *Junb* conditional deletion strategy that has since been demonstrated to cause cell-extrinsic defects in Treg development (18). Because of the predicted role of JunB in tissue-specific Tregs and previous data demonstrating broad and important roles for JunB in Th17 cells, we therefore chose to directly investigate whether JunB was important for adaptation of Treg effector function to non-lymphoid tissues.

Here, we describe a novel role for JunB in regulating the transcriptional programming of intestinal Tregs. We observed that JunB expression was highly enriched in nearly all Tregs from the intestinal lamina propria, and that Treg-specific ablation of JunB elicited an immune dysregulatory phenotype preferentially affecting the colon. In contrast to the roles of related AP-1 family TFs in Tregs, JunB was not required for the differentiation of eTregs, nor for the differentiation of tissue-specific Treg subsets in the colon or adipose tissue. Rather, we found that JunB was required for the maintenance of CD25^-^ Tregs – which included Tfr cells – leading to a loss of these populations in the spleen and Peyer’s patches (PP) of mice with Treg-restricted deletion of JunB. In the colon, JunB was required to maintain normal Treg proportions but was not required for any particular Treg population. Instead, we found that JunB regulated select components of the colonic Treg transcriptome including genes for the effector molecules Granzyme A and Granzyme B. Notably, JunB was not required for expression of any other described Treg effector molecule, suggesting that loss of JunB-dependent cytolytic gene expression in Tregs was sufficient to impair normal colonic immune homeostasis. Therefore, we have identified JunB as a critical regulator of intestinal adaptation in Tregs that controls organ-specifc effector functions.

## Results

### Junb is preferentially expressed in Tregs from the intestinal mucosa

Previous studies have suggested that *Junb* expression is significantly elevated in Tregs from the colon relative to the spleen and lymph nodes (6, 8); however, it remained unclear whether elevated *Junb* expression was a feature of all colonic Tregs or was attributable to a small subset with high *Junb* expression. To this end, we analyzed publicly-available single-cell (sc)RNA-seq data comparing Tregs isolated from various lymphoid and non-lymphoid tissues to determine the cellular distribution of *Junb* mRNA expression (4). We observed that *Junb* was highly expressed in the vast majority of colonic Tregs from both colon and skin, whereas Tregs from secondary lymphoid organs were broadly lower, with a substantial fraction below the limit of detection (Figure 1A). This suggested that increased *Junb* expression might be a general feature of Tregs in non-lymphoid organs.

**Figure 1.**
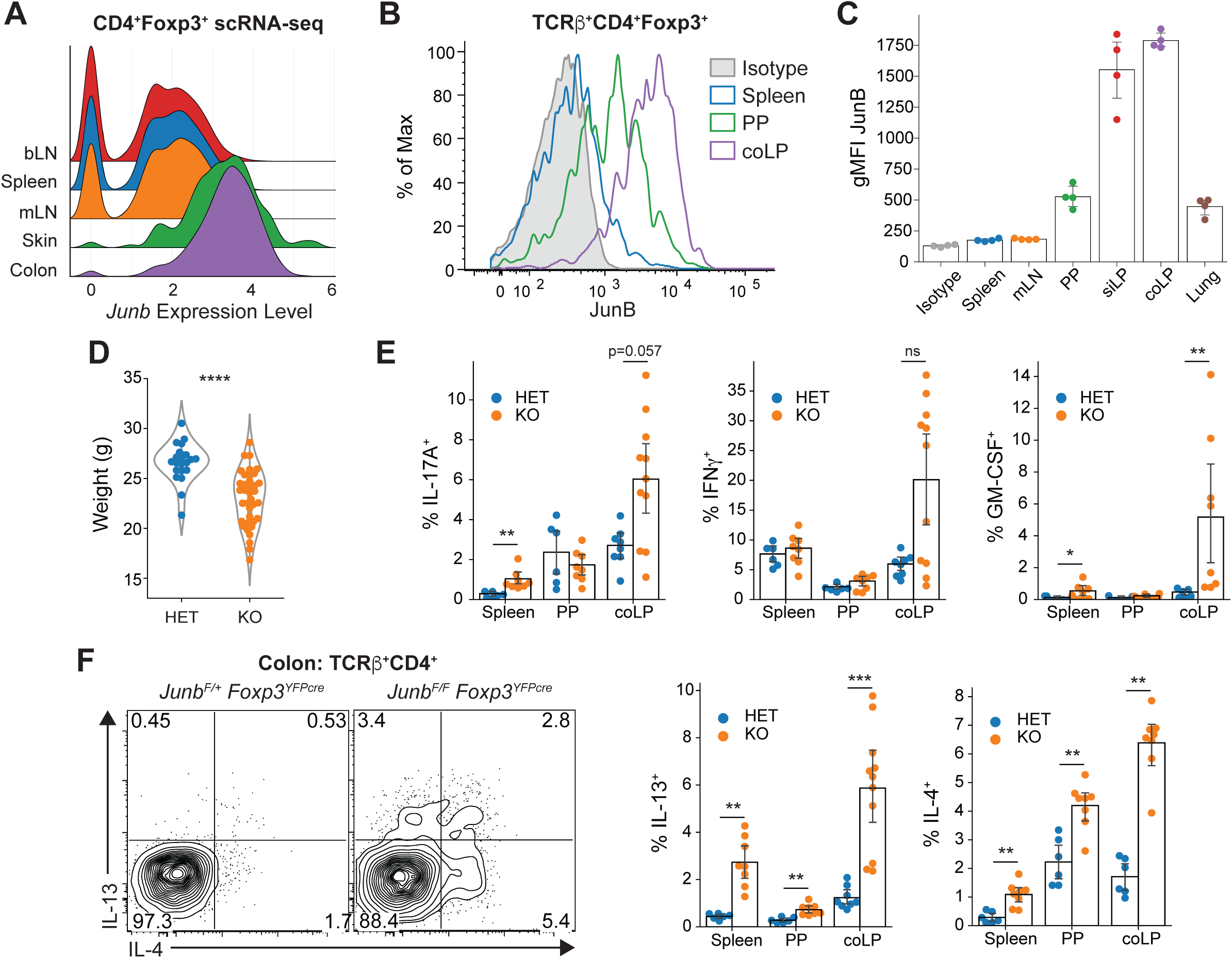
Expression of JunB in Tregs and loss of immune homeostasis following Treg-restricted deletion of *Junb.* **(A)** Expression of JunB in Treg cells across tissues determined by scRNA-seq. **(B)** Histograms and **(C)** aggregate geometric mean fluorescent intensity (gMFI) of JunB protein expression in Treg cells from indicated tissues, representative data from one of two independent experiments. **(D)** Body weights of *Junb^F/+^* (HET) or *Junb^F/F^* (KO) *Foxp3^YFPcre^* mice as determined at necropsy. Statistical significance determined using Welch’s *t*-test. **(E)** Frequencies of cytokine-producing cells among TCRβ^+^CD4^+^ cells in the indicated organs. Significance determined by Mann-Whitney U-test with Holm-Bonferroni correction. **(F)** Representative contour plots and summary data for type 2 cytokines, with pre-gating and statistics performed as in *(E)*. All results represent mean with a bootstrapped 95% confidence interval. **p < 0.01; ***p < 0.001; ****p < 0.0001; ns, not significant.

Using flow cytometry, we then determined whether JunB protein expression followed a similar pattern. In agreement with mRNA expression, JunB protein expression was substantially elevated (∼6-7-fold) in Tregs from the lamina propria of the small intestine (siLP) and colon (coLP) relative to both the spleen and mesenteric lymph nodes (mLN) (Figures 1B and 1C); however, high JunB expression was not a general feature of Tregs from all non-lymphoid tissues because Tregs isolated from the lung expressed substantially less JunB than those from the intestinal mucosa (Figure 1C). In contrast, the spleen and mLN showed a negligible increase in fluorescence intensity relative to an isotype control, suggesting that JunB protein is generally low or absent in Tregs from lymphoid organs (Figures 1B and 1C). Interestingly, Tregs from the Peyer’s Patches (PP) showed intermediate JunB expression, potentially reflecting their unique anatomy as secondary lymphoid organs embedded within mucosal tissue. Because the intestine contains almost exclusively activated eTregs, we examined whether CD62L^-^ eTregs from the spleen and PP expressed more JunB than CD62L^+^ cTregs, which have a resting phenotype. Despite the generally low expression of JunB in the spleen, we found that eTregs expressed significantly more JunB than cTregs (Supplementary Figure 1A). Additionally, we noted that JunB expression was further elevated on the PD-1^+^ subset of eTregs (Supplementary Figure 1A), which have been associated with enhanced suppressive function (19). Taken together, these observations suggested that JunB might play an important role in the function of Tregs, particularly those in the intestine.

### Ablation of JunB in Tregs impairs colonic immune homeostasis

To evaluate the requirement for JunB expression in Tregs, we generated mice with Treg-conditional ablation of *Junb* using the *Foxp3^YFP-cre^* deleter strain (20). Surprisingly, comparison of animals harboring Tregs heterozygous for *Junb* (*Junb^F/+^Foxp3^YFP-cre^*; HET) with those bearing Tregs completely deficient for *Junb* (*Junb^F/F^Foxp3^YFP-cre^*; KO) demonstrated a clear reduction in overall body weight in KO animals, suggestive of immune dysregulation. This manifestation was spontaneous and variable in severity, as indicated by the right-skewed distribution of weights (Figure 1D). However, despite weight loss, these animals did not exhibit classical signs of systemic Treg dysfunction (scaled skin, ocular irregularities, dermal thickening, splenomegaly, etc.; i.e. the “scurfy” phenotype (21)). We also observed no spontaneous mortality (up to at least 14 weeks of age; data not shown), which contrasts markedly with the ∼35% lethality in *Junb^F/F^Foxp3^YFP-cre^*mice reported by Koizumi *et al.* (*17*), suggesting that environmental differences may exert a substantial influence on JunB-dependent Treg activity. Taken together, these data suggest that JunB plays a nuanced role in Treg-mediated immune homeostasis.

Due to the biased expression of JunB in intestinal Tregs and observed weight loss in KO animals (a common manifestation of colitis), we evaluated CD4^+^ T cells for evidence of colonic cytokine dysregulation. Using flow cytometry, we found that the proportion of CD4^+^ T cells secreting the proinflammatory cytokines IL-17A, IFN*γ*, and GM-CSF were increased, with exceptionally high but variable expression in the colon of KO animals compared to controls (Figure 1E). Production of IL-17A and IFN*γ* in the colon correlated with weight deficit (Supplementary Figure 1B), suggesting that spontaneous factors – such as differences in the intestinal microbiota – were responsible for the variability in weight loss of KO animals. In contrast, we found consistently increased frequencies of CD4^+^ T cells producing the type 2 cytokines IL-4 and IL-13 in spleen, PP, and coLP (Figure 1F), demonstrating that Tregs require JunB for homeostatic regulation of CD4^+^ T cell cytokine responses in the colon.

### JunB is required to maintain organ-specific Treg proportions

To determine the Treg-intrinsic role of JunB underlying the immune dysregulation observed in KO animals, we first assessed whether JunB was required to maintain the size of the general Treg population. Intriguingly, we observed normal proportions of Tregs in the spleen, mLN, and PP of KO animals, but a substantial (nearly 2-fold) reduction in Treg frequencies in the colon (Figure 2A). However, this was not due to Treg loss *per se*, as the total numbers of Foxp3^+^ Tregs in the colon was instead increased in KO animals (Figure 2B). This observation is likely the result of inflammatory cell infiltration, because total cell numbers were selectively increased in the colon of KO animals (Supplementary Figure 2A). Therefore, JunB is required to maintain normal Treg-to-T conventional cell balance in the colon, which may be responsible for the cytokine dysregulation observed in KO mice.

**Figure 2.**
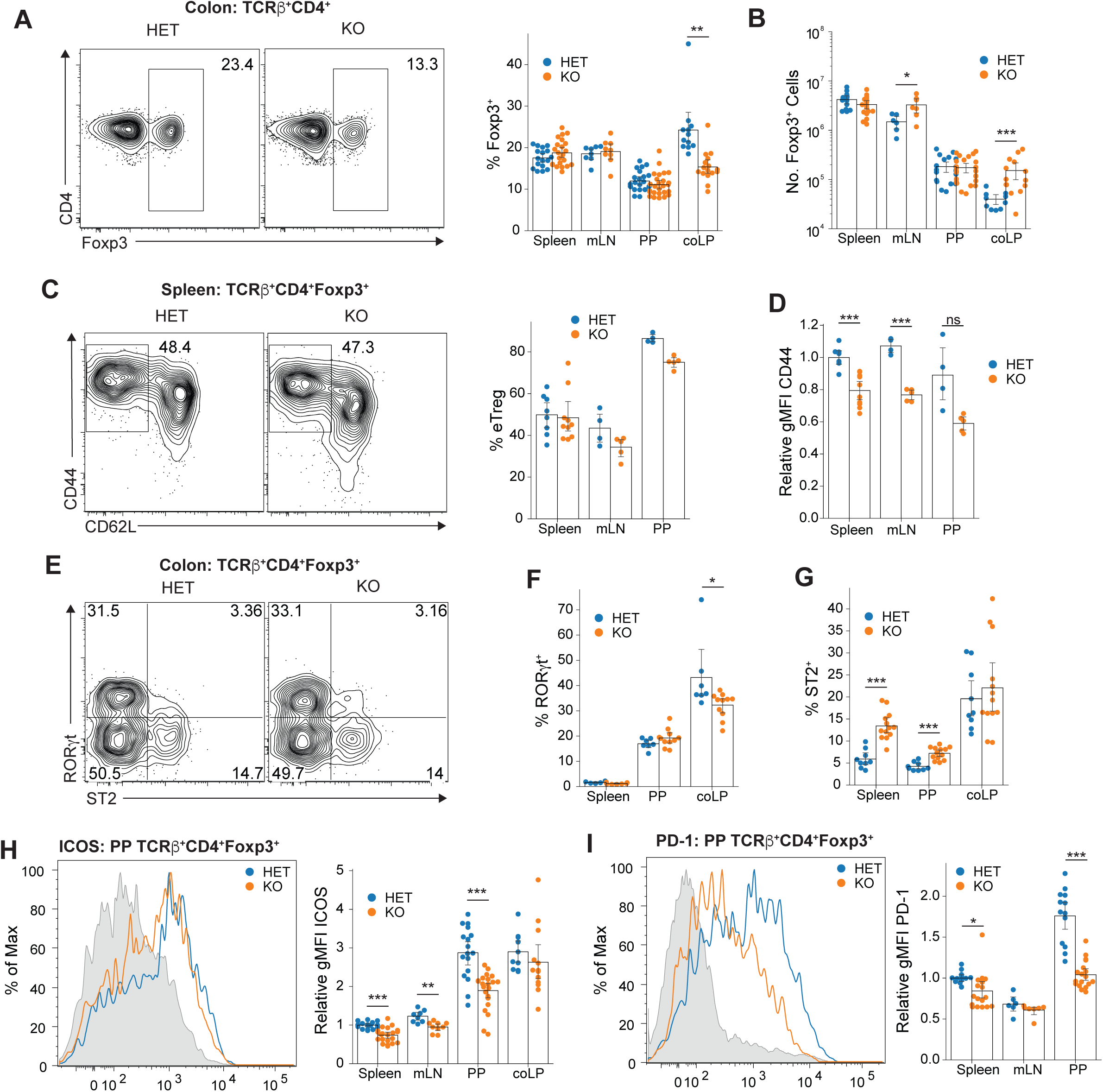
Loss of JunB causes organ-specific alterations in Treg abundance and phenotype. **(A)** Frequency of Foxp3^+^ Treg cells among TCRβ^+^CD4^+^ cells in the indicated organs. Shown are representative contour plots for the colon (left) and summary statistics for all organs (right). Statistical significance is based on a significant ANOVA interaction term followed by planned comparisons within each organ using Welch’s *t*-test with Holm-Bonferroni correction. **(B)** Number of Foxp3^+^ Treg cells within the indicated organs. Statistical analysis performed as in *(B)*, using log-transformed count values. **(C)** Representative contour plots (left) and summary data (right) showing frequency of CD44^hi^CD62L^-^ eTregs among TCRβ^+^CD4^+^Foxp3^+^ cells. No significant difference identified by Mann-Whitney U-test with Holm-Bonferroni correction. **(D)** Expression of CD44 on eTreg cells expressed as relative gMFI, normalized to the mean gMFI for splenic eTreg cells from HET animals in each individual experiment. Statistical significance determined by Welch’s *t*-test with Holm-Bonferroni correction. **(E)** Representative contour plots showing colonic RORγt^+^ and ST2^+^ Treg cells. **(F)** Frequency of ROR*γ*t^+^ and **(G)** ST2^+^ Treg cells among all Treg cells. Significance determined using Mann-Whitney U-test with Holm-Bonferroni correction. **(H)** Representative histogram (left) showing ICOS expression on Treg cells from PP of HET (blue) or KO (orange) animals, splenic Treg cells from HET animals are shown in gray for reference. Relative gMFI of ICOS, normalized as in *(D)*. Significant organ:genotype interaction by ANOVA followed by planned comparisons using Welch’s t-test with Holm-Bonferroni correction. **(I)** Representative histogram (left) showing PD-1 expression on Treg cells from PP of HET (blue) or KO (orange) animals, splenic CD4^+^Foxp3^-^ T cells are shown in gray for reference. Relative gMFI (right) of PD-1 expression on Foxp3^+^ Treg cells, normalized as in *(D)* with statistical significance determined as in *(H)*. All summary data represent mean with a bootstrapped 95% confidence interval. *p < 0.05; **p < 0.01; ***p < 0.001; ****p < 0.0001; ns, not significant.

### eTreg differentiation does not depend on JunB

The vast majority of Tregs in the colon are highly active eTreg cells (CD44^hi^CD62L^-^) whereas the spleen and LN contain substantial populations of both eTregs and cTregs (CD44^lo^CD62L^+^) (1). Because JunB expression was elevated in eTregs (Supplementary Figure 1B), we investigated the possibility that JunB was selectively required to maintain the eTreg population. However, KO animals showed only a relatively small decrease in eTregs frequencies in the spleen, mLN, and PP that was not statistically significant, indicating that maintenance of eTregs does not generally depend on JunB (Figure 2C). This notion was further supported by an increase in the frequency of Tregs expressing KLRG1, which has been considered a preferential marker of eTregs (9) (Supplementary Figure 2B). Interestingly, despite a negligible decrease in the frequency of eTregs, we did observe a reduction in the expression level of CD44 on eTregs lacking JunB (Figures 2C and 2D). Nevertheless, the expression of CD44 on eTregs from KO animals was still much higher than that of cTregs (Figure 2C and Supplementary Figure 2C), which suggested that CD44 expression was not strictly JunB-dependent and was unlikely to be biologically significant.

### Differentiation of specialized colonic Treg subpopulations does not require JunB

Colonic Tregs can be subdivided into two primary populations with distinct antigen specificities and transcriptional programs: ROR*γ*t^+^ Tregs and GATA-3^+^/ST2^+^ Tregs (3). Therefore, we considered that the decrease in colonic Treg frequencies in KO mice might reflect selective loss of one of these populations. Flow cytometric analysis revealed only a small reduction in ROR*γ*t^+^ Tregs in the colon, whereas the same population in the spleen and PP was unaffected (Figure 2E). However, the ST2^+^ Treg population was largely unchanged in the colon but was significantly increased in the spleen and PP of KO mice (Figures 2E and G). Interestingly, the size of the ST2^+^ Treg population was correlated with the magnitude of cytokine production (Supplementary Figure 2D), suggesting that this reflected an ongoing Treg response to inflammation in KO animals and that Tregs remain responsive to inflammatory signals in the absence of JunB. Consistent with similar frequencies of ROR*γ*t^+^ Tregs, which are Helios^-^ (^8^), we observed no change in the ratio of Helios^+^ to Helios^-^ Tregs in KO mice, suggesting that JunB was not preferentially required for the maintenance of either thymus-or peripherally-derived Tregs, respectively (Supplementary Figure 2E) (3). Altogether, these data suggest that JunB plays a subset-independent, organ-specific role in Treg function.

### Reduced expression of ICOS and PD-1 on Tregs in the absence of JunB

Because we observed decreased expression of CD44 in JunB-deficient Tregs, we next examined the expression of additional activation markers. Inducible costimulator (ICOS) and PD-1 are preferentially expressed on eTregs and have been previously implicated in both the population maintenance and suppressive function of Tregs (22–26). Furthermore, we observed that JunB expression was significantly elevated on the PD-1^hi^ subset of eTregs in both spleen and PP (Supplementary Figure 1B), suggesting that JunB might be particularly important for the function of highly-activated eTregs. Using flow cytometry we observed a significant reduction of both ICOS and PD-1 expression in KO mice, most notably in the PP (Figures 2H and 2I). However, KO animals still exhibited increased ICOS expression in the PP and coLP relative to spleen and mLN, suggesting that ICOS expression is not completely dependent on JunB (Figure 2H). Taken together, these data suggested that JunB might be important in driving expression of ICOS and PD-1 or, alternatively, in maintaining a subpopulation of activated Tregs that exhibit high ICOS and PD-1 expression.

### Tfr cells are critically dependent on JunB

Tfr cells are a subset of Tregs which localize to germinal centers (GCs) and are thought to interact with both T follicular helper (Tfh) cells and GC B cells to control the processes of affinity maturation and class switch recombination (CSR) (2). Considering that high expression of ICOS and PD-1 are defining features of follicular T cells (both Tfr and Tfh), we hypothesized that reduced expression of ICOS and PD-1 in KO animals might reflect loss of Tfr cells, which are particularly abundant in the PP. Indeed, we observed a striking near-complete loss of Bcl6^+^ PD-1^hi^ Tfr cells in both the spleen and PP of KO mice, indicating that JunB is essential for the development or maintenance of the Tfr population (Figure 3A). The loss of Tfr cells did not lead to altered proportions of Tfh cells or GC B cells in the spleen and PP of KO animals, consistent with previous studies directly ablating Tfr cells (27) (Supplementary Figures 3A and 3B). However, we did observe a trend towards increased IgG1 CSR among GC B cells and a significant elevation of CD38^+^IgG1^+^ memory B cells in both the spleen and PP (Supplementary Figures 3C and D). This is consistent with enhanced production of IL-4 – an established driver of IgG1 CSR – in KO animals (Figure 1F). Furthermore, we observed increased frequencies of IgA-bound microbes in the feces of KO mice, suggesting that secretory IgA was increased in magnitude or altered in specificity (Supplementary Figure 3E). Altogether, JunB is a critical regulator of Tfr cell differentiation or maintenance and is required for normal steady-state antibody responses, particularly in the intestine.

**Figure 3.**
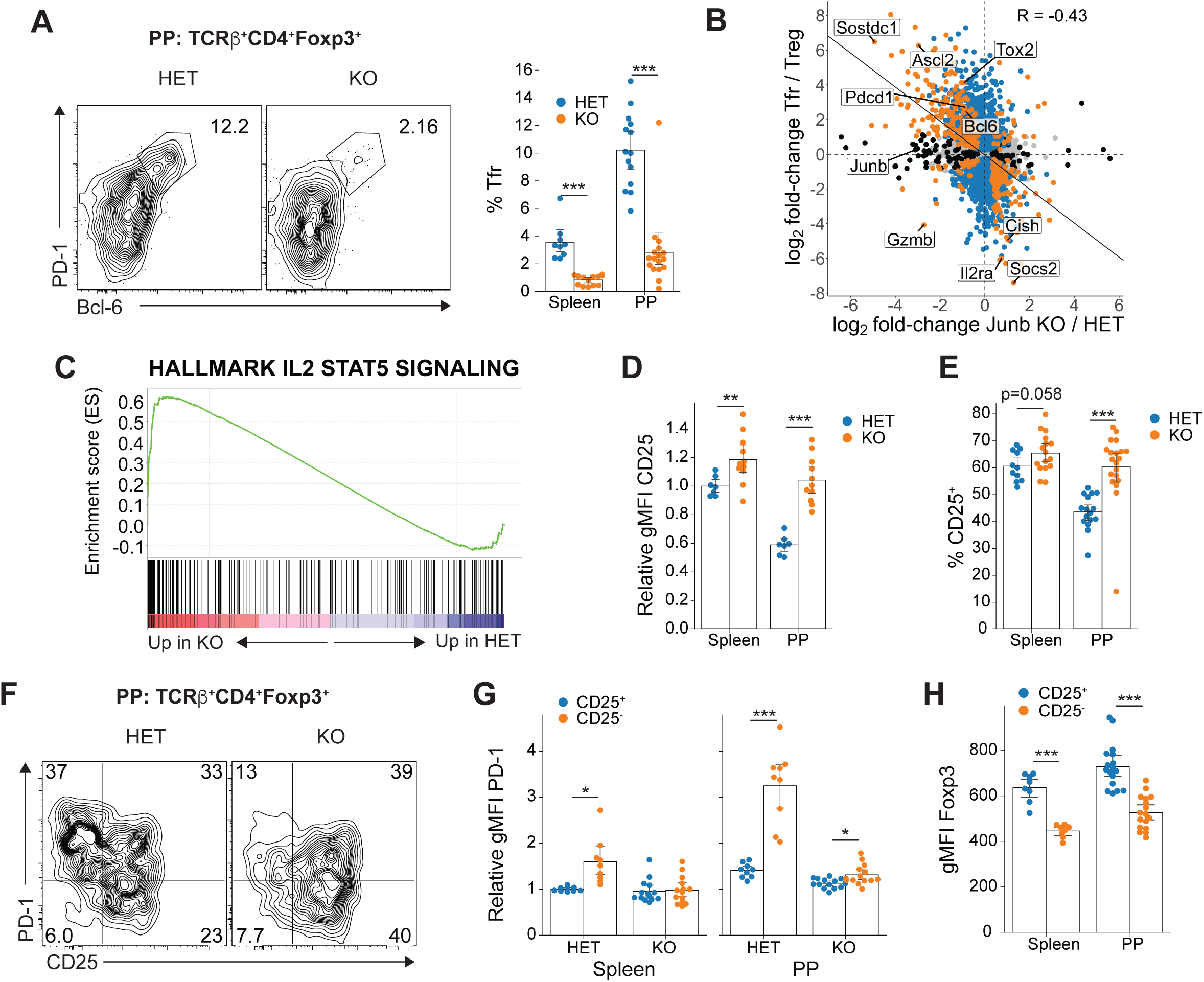
Loss of JunB selectively ablates Tfr cells and CD25^-^ Treg cells. **(A)** Representative flow cytometry plots showing PD-1^hi^Bcl-6^+^ Tfr cells (left). Summary data (right) are compiled from Tfr cells identified as either PD-1^hi^Bcl-6^+^ or PD-1^hi^CXCR5^+^. **(B)** Relationship between the Tfr cell transcriptome and JunB-dependent transcriptome. Orange indicates significant DE in both datasets, while blue and black indicate significant DE in Tfr and JunB datasets, respectively. A linear model was fitted to points with significant DE in at least one of the two datasets; a significant correlation was detected using the Pearson method (p < 2×10^-16^). **(C)** GSEA enrichment plot for the MSigDB Hallmarks IL-2 / STAT5 signaling gene set in JunB HET vs KO RNA-seq. **(D)** Relative expression of CD25 on TCRβ^+^CD4^+^Foxp3^+^ cells (normalized to the mean gMFI for splenic Foxp3^+^ cells in HET mice from each experiment) and **(E)** percentage of CD25^+^ cells as determined by flow cytometry. Significance determined by Welch’s *t*-test with Holm-Bonferroni correction. **(F)** Representative contour plot showing the relationship between CD25 and PD-1 expression. **(G)** Relative expression of PD-1 on CD25^+^ and CD25^-^ Treg cell subsets. Significance determined by 3-way ANOVA followed by Welch’s *t*-test with Holm-Bonferroni correction. **(H)** Expression of Foxp3 on CD25^+^ and CD25-subsets of TCRβ^+^CD4^+^Foxp3^+^ cells as determined by flow cytometry. Data are pooled from both HET and KO animals because ANOVA indicated no significant effect of genotype. Statistical significance determined by Welch’s *t*-test with Holm-Bonferroni correction. All summary data represent mean with a bootstrapped 95% confidence interval. *p < 0.05; **p < 0.01; ***p < 0.001.

To uncover the mechanism of JunB function in PP Tregs, we next assessed JunB-dependent transcriptional changes using RNA-sequencing (RNA-seq). We observed reduced expression of numerous canonical Tfr/Tfh genes, including *Pdcd1* (PD-1), *Bcl6*, and *Ascl2* (Supplementary Figure 3F), consistent with loss of Tfr cells. While Tfr constitute only 10% of PP Tregs, their complete loss in KO animals complicated the extraction of candidate JunB targets from this dataset. To this end, we compared publicly available data defining the Tfr-specific transcriptome with JunB-dependent genes in PP Tregs to identify shared and divergent transcripts (28). Unsurprisingly, we observed a strong correlation between JunB-dependent and Tfr-associated transcripts, confirming that JunB is essential for Tfr cells and that the majority of differentially-expressed (DE) genes reflected complete loss of the Tfr transcriptome (Figure 3B). However, a small number of genes were found to be JunB-dependent but not correlated with the Tfr cells, such as *Gzmb* (Granzyme B), which has been implicated as a suppressive mechanism employed by Tregs (29–31). Importantly, despite the significant decrease in ICOS expression observed in KO PP Tregs, we found no evidence of reduced *Icos* transcription by RNA-seq (Supplementary Figure 3G), indicating that *Icos* is not a direct transcriptional target of JunB in Tregs. Taken together, these data demonstrate a novel and essential role for JunB in Tfr cells.

### JunB is essential for maintenance of CD25^-^ Tregs

To determine whether the essential role for JunB in Tregs might be due to regulation of a critical signaling pathway or cellular function, we performed gene-set enrichment analysis (GSEA) on the JunB-dependent transcriptome. Surprisingly, the dominant gene signature was an increase in the IL-2/STAT5 signaling pathway, with leading-edge genes including the SOCS-family members *Cish* and *Socs2*, as well as the IL-2 receptor *α* chain (*Il2ra;* CD25) (Figures 3B and 3C). This was particularly intriguing because – despite the long-recognized importance of IL-2 signaling for the development, maintenance, and function of Tregs – recent evidence has established that Tfr cells are uniquely IL-2-independent, express low levels of CD25, and are inhibited by high levels of IL-2 (28, 32, 33). Assessment of CD25 expression revealed a striking elevation in both the level of CD25 protein and the percentage of CD25^+^ Tregs in KO mice (Figures 3D and 3E), demonstrating a marked loss of the CD25^-^ Treg population that encompasses Tfr cells. Because we observed inflammatory cytokine dysregulation in KO mice, we considered that CD25 upregulation could be driven by STAT5 signaling resulting from increased IL-2 production by activated CD4^+^ T cells. However, we failed to observe an increase in IL-2-expressing T cells in the spleen or PP (Supplementary Figure 3H), suggesting that this was not a contributor to the loss of CD25^-^ Tregs. Importantly, we noted that CD25^-^ Tregs displayed elevated expression of both PD-1 (Figures 3F and 3G) and ICOS (Supplementary Figures 3I and 3J), indicating that reduced ICOS and PD-1 expression on KO Tregs reflected loss of the CD25^-^ Treg population. Overall, these data demonstrate a critical role for JunB in the maintenance of CD25^-^ Tregs and, consequently, Tfr cells.

### Treg lineage stability is not influenced by JunB

Next, we sought to explain the unique dependence of CD25^-^ Tregs on JunB. Consistent with previous reports (32, 34), we noted that CD25^-^ Tregs displayed reduced expression of Foxp3, the lineage-defining TF for Tregs (Figure 3H). Considering our previous work demonstrating that Th17 cell identity is JunB-dependent, we reasoned that low Foxp3 expression might render CD25^-^ Tregs particularly prone to lineage instability (i.e. loss of Foxp3), which was previously shown for PP Tregs (35) and could potentially result in their conversion to pathogenic “ex-Treg” effectors. To this end, we attempted a lineage-tracing experiment by crossing *Junb Foxp3^YFP-cre^* mice to those bearing a *Rosa26^STOP-ZsGreen^* allele (36) such that cells that have expressed *Foxp3* are permanently labeled with ZsGreen. To our surprise, the vast majority of animals born from these crosses developed visibly green skin and internal organs, indicating germline recombination at the *Rosa* locus (data not shown). In the small number of animals obtained that were not visibly green, we identified widespread promiscuous Cre activity affecting ∼10-30% of all hematopoietic lineages, with further increases in CD4^+^ Foxp3^-^ cells (Supplementary Figure 4A). Additionally, we identified deleted *Junb* alleles in Cre^-^ offspring from *Foxp3^YFP-cre^*breedings (data not shown). Overall, this indicated that *Junb* was spontaneously deleted in the germline of *Foxp3^YFPcre^*-bearing dams and is consistent with several previous reports indicating promiscuous activation of *Foxp3*-driven Cre recombinase, including *Foxp3^YFPcre^*^(37, 38^).

To circumvent these complications, we bred *Junb* floxed mice to *Foxp3^EGFP-cre-ERT2^* (36) and Rosa26*^STOP-ZsGreen^* mice to generate Junb*^F/+^* Foxp3*^EGFP-cre-ERT2^* Rosa26*^STOP-ZsGreen^* (iHET) and *Junb^F/F^ Foxp3^EGFP-cre-ERT2^ Rosa26^STOP-ZsGreen^* (iKO) mice which delete *Junb* and activate ZsGreen expression in Foxp3-expressing cells only in the presence of tamoxifen. We then evaluated Treg stability by assessing the proportion of TCR*β*^+^CD4^+^ZsGreen^+^ cells that lost Foxp3 expression 14 days after administration of tamoxifen by oral gavage. Despite robust labeling efficiency (Supplementary Figure 4B), we observed no significant change in the proportion of ZsGreen^+^ cells that lost Foxp3 expression in treated iKO versus iHET mice (Figure 4A), indicating that JunB is not required for Treg lineage stability. While we did observe a small increase in Foxp3^-^ cells among ZsGreen^+^ cells in the colon, this change was not statistically significant and likely reflects enrichment of the normally present ex-Treg population due to the decrease in Treg frequencies in this organ (Figure 4B). Importantly, JunB depletion did not affect the frequency of ex-Tregs in the PP (Figure 4A) – which normally hosts a large CD25^-^ Treg population – confirming that loss of CD25^-^ Tregs in the absence of JunB is not due to Treg lineage instability.

**Figure 4.**
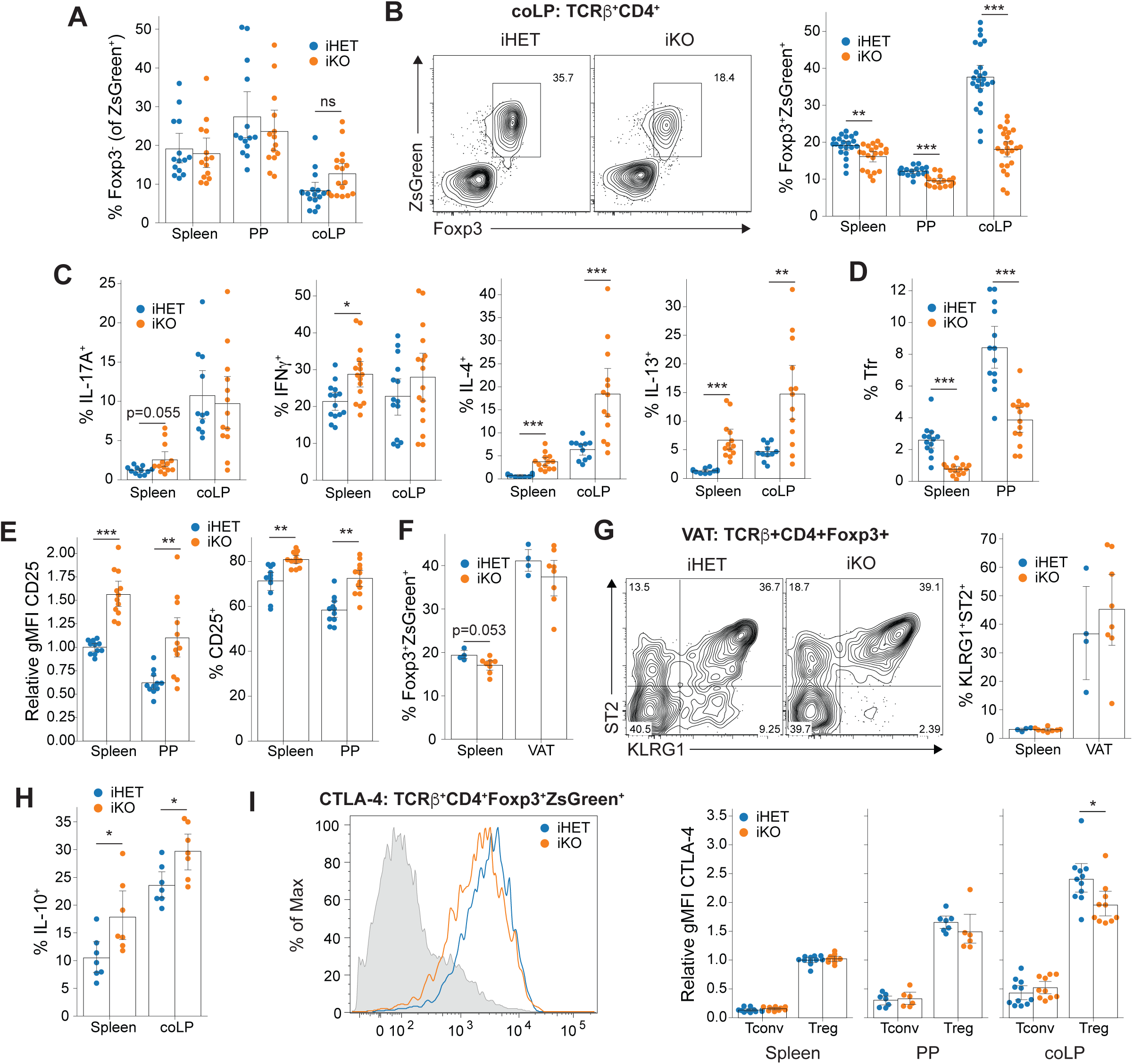
JunB does not influence Treg stability but is continuously required for select Treg functions. **(A)** Treg cell stability, determined as frequency of Foxp3^-^ cells among labeled TCRβ^+^CD4^+^ZsGreen^+^ cells 14 days after tamoxifen-induced *Junb* deletion. Statistical significance determined by Mann-Whitney U-test with Holm-Bonferroni correction. **(B)** Representative flow cytometry plots (left) and summary statistics (right) showing frequency of Foxp3^+^ZsGreen^+^ Treg cells among TCRβ^+^CD4^+^ cells following tamoxifen-induced deletion of *Junb*. Statistical testing by two-way ANOVA identified a significant organ:genotype interaction and was followed by Welch’s *t*-test with Holm-Bonferroni correction. **(C)** Frequency of TCRβ^+^CD4^+^ZsGreen^-^ cells expressing the indicated cytokines 14 days after tamoxifen administration. Significance determined by Mann-Whitney U-test with Holm-Bonferroni correction. **(D)** Frequency of PD-1^hi^Bcl-6^+^ Tfr cells among TCRβ^+^CD4^+^Foxp3^+^ZsGreen^+^ cells. Statistical significance determined by Mann-Whitney U-test with Holm-Bonferroni correction. **(E)** Relative expression of CD25 (left) and frequency of CD25^+^ cells (right) among TCRβ^+^CD4^+^Foxp3^+^ZsGreen^+^ Treg cells. Statistical significance determined by Mann-Whitney U-test with Holm-Bonferroni correction. **(F)** Frequency of Foxp3^+^ZsGreen^+^ Treg cells among TCRβ^+^CD4^+^ cells in the spleen and VAT of 14-16 week old male mice. Statistical significance determined by Mann-Whitney U-test with Holm-Bonferroni correction. **(G)** Representative contour plots (left) and summary statistics (right) showing the frequency of TCRβ^+^CD4^+^Foxp3^+^ZsGreen^+^ displaying the characteristic VAT Treg phenotype. **(H)** Frequency of TCRβ^+^CD4^+^ZsGreen^+^ cells expressing IL-10. Statistical significance determined by Mann-Whitney U-test with Holm-Bonferroni correction. **(I)** Representative histogram showing CTLA-4 expression in Tregs from iHET and iKO mice compared to splenic CD4^+^Foxp3^-^ cells in gray (left) and summary data (right) showing expression of CTLA-4 as determined by flow cytometry, normalized to the gMFI of splenic Treg cells from iHET mice for each experiment. Statistical significance determined by Mann-Whitney U-test with Holm-Bonferroni correction. All summary data represent mean with a bootstrapped 95% confidence interval. *p < 0.05; **p < 0.01; ***p < 0.001.

### Continuous expression of JunB is required to maintain Treg homeostasis

Given that JunB plays important roles in conventional CD4^+^ T cells and that homozygous germline deletion of *Junb* results in embryonic lethality through non-immunological mechanisms (39), we considered that our previous findings using *Foxp3^YFPcre^*might be confounded by promiscuous deletion of *Junb*. Therefore, we sought to confirm these observations using tamoxifen-induced deletion in iHET and iKO mice. This model system also allowed us to evaluate whether the requirement for JunB in Tregs was continuous or transient, because *Junb* was deleted in mature Tregs rather than during initial Treg differentiation. Surprisingly, only two weeks after initiating deletion of *Junb*, we observed complete recapitulation of the colon-specific JunB-dependent defect in Treg proportions (Figure 4B); however, we also observed a small but statistically-significant reduction in Treg frequencies in the spleen and PP (Figure 4B), suggesting that time-dependent compensatory mechanisms may have obscured this difference in *Foxp3^YFPcre^* mice. Furthermore, unlike the *Foxp3^YFPcre^*phenotype, we observed that numbers of both Tregs (Supplementary Figure 4C) and total cells (Supplementary Figure 4D) were the same in iHET and iKO animals, rather than increased, the latter suggesting that inflammatory infiltration was less pronounced at short intervals following deletion. These data confirm that JunB is essential to maintain normal colonic Treg proportions, likely by facilitating suppression of inflammatory T cell accumulation.

As in KO *Foxp3^YFPcre^* mice, we observed a consistent elevation in IL-4 and IL-13 expression among CD4^+^ T cells in iKO mice, with variable expression of both IL-17A and IFN*γ* (Figure 4C). Similarly, eTreg frequencies and CD44 expression were also reduced to a comparable degree (Supplementary Figure 4E). Although iKO mice displayed a small decrease in ROR*γ*t^+^ Tregs in the colon (Supplementary Figure 4F), we no longer observed increased ST2^+^ Treg frequencies in the spleen and PP (Supplementary Figure 4G), suggesting that this was a response to prolonged inflammation in KO *Foxp3^YFPcre^* mice. Inducible *Junb* deletion recapitulated the major reduction in Tfr cell frequencies (Figure 4D) and increased CD25 expression (Figure 4E), again despite no increase in IL-2 expression by CD4^+^ T cells in iKO compared to iHET mice (Supplementary Figure 4H). Interestingly, we did not observe a change in ICOS expression in the spleens of iKO mice – only in the PP – further supporting that *Icos* is not a direct transcriptional target of JunB (Supplementary Figure 4I). Although we failed to observe a significant decrease in PD-1 expression in iKO mice, there did appear to be a trend towards reduced expression (Supplementary Figure 4J). Overall, these data confirm a consistent and continuous role for JunB in the maintenance of colonic T cell homeostasis, Treg abundance, CD25^-^ Tregs and Tfr cells.

### JunB is not required for adipose-resident Tregs

Tregs that reside in the VAT are among the best-studied tissue-resident Treg populations. Previous data have shown that VAT Tregs require the transcription factors BATF and IRF4 (9)– two major binding partners of JunB in CD4^+^ T cells (40–42) – suggesting that JunB could be important in VAT Tregs. Furthermore, the loss of tissue-resident Treg homeostasis in the colon suggested that JunB might be generally important for tissue-resident Tregs; however, the reduced weight of KO *Foxp3^YFP-cre^* mice resulted in animals with abnormally low body fat, precluding the isolation of VAT lymphocytes. To circumvent this limitation, we administered tamoxifen to healthy 14-16-week-old male iHET and iKO mice and analyzed VAT-resident Tregs 14 days later. Despite consistent impairment of the colonic Treg population using this model, we observed no apparent change in the frequency of Tregs within the VAT of iKO mice (Figure 4F). Furthermore, despite a high degree of variability in the data, it was clear that JunB was not required to maintain the characteristic KLRG1^+^ST2^+^ VAT Treg phenotype (Figure 4G). Therefore, the requirement for JunB in tissue homeostasis appears to be organ-specific.

### Expression of canonical suppressive genes is largely JunB-independent

Altogether, the data strongly implicated JunB in the selective maintenance of Treg-mediated immune homeostasis in the intestine; however, the transcriptional targets of JunB underlying this requirement remained unclear. Tregs employ several distinct mechanisms to negatively-regulate immune responses, including competitive consumption of IL-2 (via CD25), secretion of anti-inflammatory cytokines (e.g. IL-10, IL-35, TGF-*β*), contact-dependent inhibition of antigen presenting cell (APC) function (e.g. CTLA-4, LAG-3), production of immunosuppressive adenosine (CD39 and CD73), and direct cytolysis of APCs and effector T cells (e.g. Granzyme A/B) (29-31, 43, 44). Because of the strong increase in CD25 expression, we considered defective IL-2 consumption unlikely to explain the JunB-dependent defect in Treg function; however, alternative mechanisms remained to be explored. Further analysis of the PP RNA-seq dataset suggested that the majority of Treg suppressive genes were not JunB-dependent, including both secreted [IL-10, IL-35 (*Ebi3*), TGF-*β*] and membrane-bound mediators [CTLA-4, CD39 (*Entpd1*), CD73 (*Nt5e*), and LAG-3] (Supplementary Figure 4K). Interestingly, Granzyme A (*Gzma*) and Granzyme B (*Gzmb*) were both highly expressed and highly JunB-dependent suggesting a potential role for JunB in facilitating suppression through Treg-mediated cytolysis (Figure 3B, and Supplementary Figures 3F and 4K).

Because IL-10 and CTLA-4 are among the most important and well-explored suppressive mechanisms used by intestinal Tregs (3), we further evaluated their expression by flow cytometry in tamoxifen-treated iHET and iKO mice. Aligned with results from RNA-seq, we observed a small increase in the frequency of Tregs expressing IL-10, confirming that JunB is not required for *Il10* expression and that JunB-deficient Tregs were able to respond to altered homeostasis by upregulating select suppressive functions (Figure 4H and Supplementary Figure 4K). However, despite no change in *Ctla4* mRNA expression by RNA-seq (Supplementary Figure 4K), we did observe a small decrease in total CTLA-4 protein in Tregs from iKO animals, suggesting JunB may influence CTLA-4 expression indirectly (Figure 4I). Nevertheless, *Junb*-deficient Tregs still expressed large amounts of CTLA-4 relative to non-Tregs (Figure 4I), and neither JunB-KO nor JunB-iKO mice showed evidence of the fatal lymphoproliferative disease that occurs following Treg-specific ablation of CTLA-4 (45). This suggested that reduced CTLA-4 expression was a minimal contributor to the immune dysregulation following JunB ablation in Tregs. Taken together, these results demonstrate that the majority of Treg effector molecules show JunB-independent expression, whereas *Gzma* and *Gzmb* appear to be highly JunB-dependent.

### JunB controls expression of metabolic genes in CD25^-^ PP Tregs

Despite extensive characterization of the cellular phenotype resulting from Treg-specific JunB deficiency, the direct transcriptional targets of JunB underlying these effects remained unclear. A major challenge in distinguishing direct transcriptional targets from collateral effects was the dominant influence of Tfr cell loss on the PP RNA-seq data, as well as likely confounding effects of the ongoing inflammatory response. We reasoned that short-term, inducible deletion of JunB using the *Foxp3^EGFP-cre-ERT2^*system would circumvent both population loss and inflammation-induced transcriptional changes, thereby allowing us to identify direct transcriptional targets of JunB using RNA-seq. Although we observed reduced colonic Treg frequencies and aberrant cytokine responses 14 days after tamoxifen-induced JunB deletion (Figures 4B and 4C), by 7 days post-tamoxifen administration we noted that colonic Treg proportions appeared normal, despite complete absence of JunB protein (Supplementary Figs 5A and 5B). Therefore, we sorted the JunB-dependent CD25^-^TCR*β*^+^CD4^+^*Foxp3*^EGFP+tdTomato+^ Treg population from the PP of *Junb^F/+^ Foxp3^EGFP-cre-ERT2^ Rosa26^STOP-tdTomato^* or *Junb^F/F^ Foxp3^EGFP-cre-ERT2^ Rosa26^STOP-tdTomato^* mice 7 days after tamoxifen administration for analysis by RNA-seq. Surprisingly, our approach effectively reduced the number of DE genes from over 800 in the *Foxp3^YFP-cre^* PP dataset to just 44 in CD25^-^ PP Tregs with particular sensitivity to JunB (FDR < 0.05; Figure 5A).

**Figure 5.**
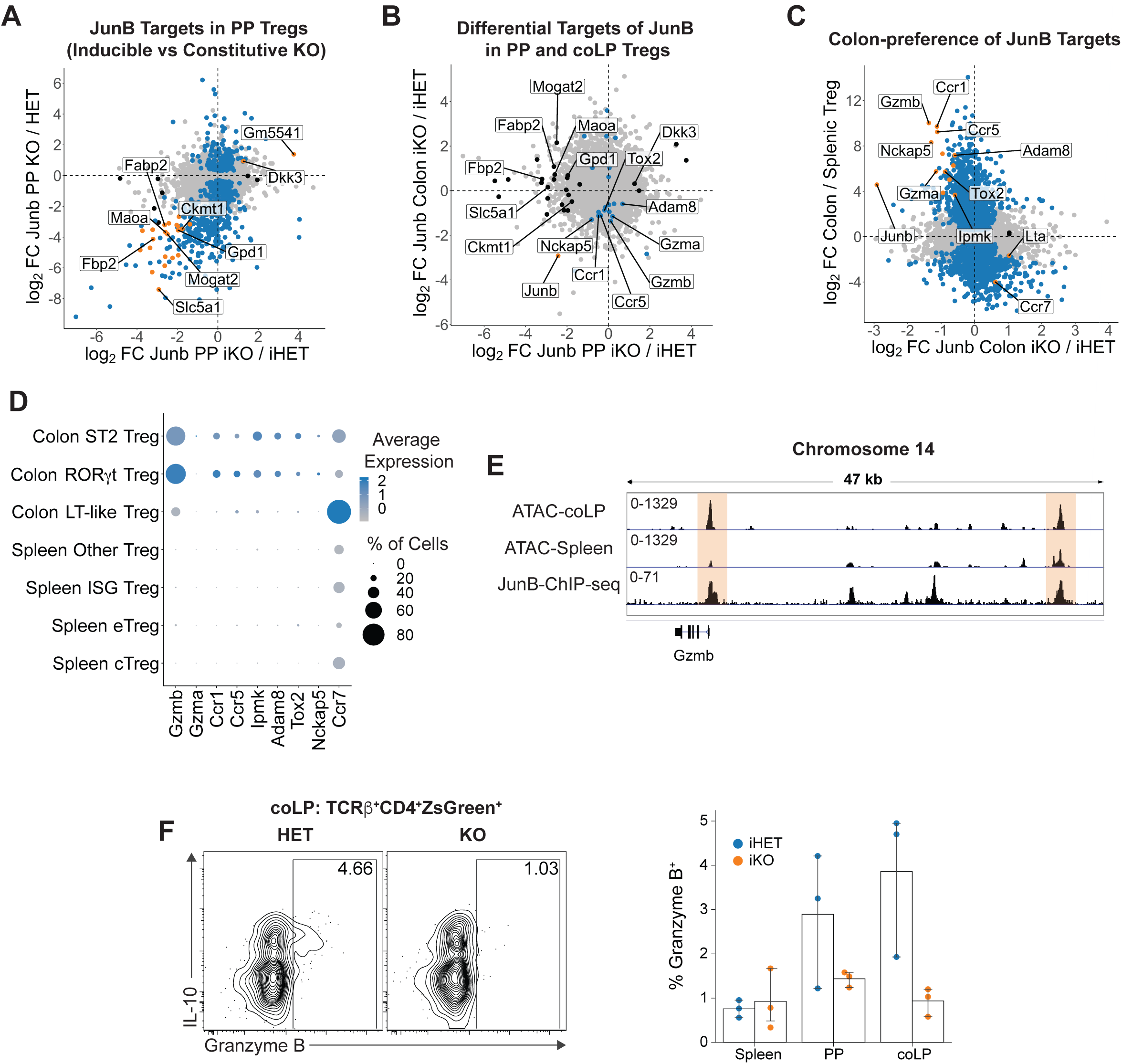
Control of the colonic Treg gene expression program by JunB. **(A)** Comparison of DE in CD25^-^ PP Treg from iHET vs iKO mice (x-axis) and DE in PP Tregs from HET vs KO mice (y-axis). Genes are colored by significance (FDR < 0.05) in both datasets (orange), only iKO vs iHET Tregs (black), or only HET vs KO Tregs (blue). **(B)** Comparison of DE in CD25^-^ PP Treg (x-axis) vs. colonic Tregs (y-axis) from iHET vs iKO mice. Genes are colored by significance (FDR < 0.05) in both datasets (orange), only JunB-iKO/iHET CD25^-^ PP Tregs (black), or only JunB-iKO/iHET colonic Tregs (blue). **(C)** Comparison of DE in colonic Tregs from iHET vs iKO mice (x-axis) with Immgen DE comparing colonic vs. splenic Tregs (y-axis). Genes are colored by significance (FDR < 0.05) in both datasets (orange), only JunB-iKO/iHET colonic Tregs (black), or only colonic/splenic Tregs (blue). **(D)** Dot plot summarizing scRNA-seq data showing both average expression levels (color) and frequency of cells (size) expressing JunB-dependent genes among the indicated splenic and colonic Treg populations. Cells identities were assigned based on the expression of canonical markers or sets of related genes [(LT; lymphoid tissue), (ISG; interferon-stimulated genes)]. **(E)** Chromatin accessibility and binding of JunB at the *Gzmb* locus. ATAC-seq data shows RPKM-normalized signal values for colonic or splenic Tregs; ChIP-seq shows RPKM-normalized signal for JunB binding in cultured Th17 cells. **(F)** Expression of Granzyme B in colonic Tregs. Representative contour plots gated on colonic Tregs (left) and summary data (right) from one experiment. All summary data represent mean with a bootstrapped 95% confidence interval.

Comparison of the JunB-dependent transcripts identified in total PP Tregs from *Foxp3^YFP-cre^* mice with those identified following short-term inducible *Junb* deletion showed a coordinated trend towards reduced expression of many genes, despite only a portion of them reaching statistical significance (Figure 5A). Interestingly, of the genes significantly DE in both datasets, we observed that many of them were related to metabolism – though with unexplored relevance in T lymphocytes – suggesting that JunB might be important for maintaining metabolic fitness in CD25^-^ Treg cells (Figure 5A). This was further supported by GSEA, which identified 4 metabolic pathways as significantly underrepresented in JunB-deficient PP Tregs (Supplementary Figure 5C). Notably, these included oxidative and fatty acid metabolism, which are unusually important in Tregs relative to other CD4^+^ T cells (46). Interestingly, GSEA of RNA-seq data comparing Tfr and Treg cells (28) demonstrated that both oxidative and glycolytic metabolism were among the top pathways distinguishing these cells, suggesting that Tfr cells are highly metabolic relative to other Tregs and may therefore be uniquely susceptible to metabolic impairment (Supplementary Figure 5D). These data are aligned with reports demonstrating that the PI3K-Akt-mTORC1 pathway – a critical regulator of growth and energy homeostasis – is particularly important for differentiation of Tfr cells (47, 48). Therefore, JunB regulates a subset of metabolic genes in CD25^-^ PP Tregs, suggesting that impaired metabolic fitness may explain loss of these cells following JunB deletion.

### JunB regulates a colonic gene expression program distinct from that of PP Tregs

To identify JunB-dependent transcripts in colonic Tregs, we followed a similar approach as for PP Tregs. Seven days after tamoxifen administration, TCR*β*^+^CD4^+^*Foxp3*^EGFP+tdTomato+^ Tregs were sorted from the colons of *Junb^F/+^ Foxp3^EGFP-cre-ERT2^* Rosa26*^STOP-tdTomato^* or Junb*^F/F^*Foxp3*^EGFP-cre-ERT2^* Rosa26*^STOP-tdTomato^* mice for RNA-seq analysis. Similar to what we observed for the PP, short-term deletion of JunB identified just 26 genes with significant DE in colonic Tregs, suggesting that these transcripts exhibit strong dependence on JunB (Supplementary Figure 5E). Surprisingly, we observed no shared JunB-dependent genes when comparing coLP Treg to CD25^-^ PP Treg RNA-seq from *Foxp3^EGFP-cre-ERT2^* mice (Figure 5B), indicating that JunB plays distinct roles in CD25^-^ PP Tregs and colonic Tregs.

We assessed the contribution of JunB to colonic Treg programming by comparing JunB-dependent transcripts in colonic Tregs with those enriched in colonic relative to splenic Tregs using data from the Immgen consortium (49). Interestingly, this revealed that essentially all JunB-dependent genes were preferentially expressed in colonic Tregs, suggesting that JunB is an important regulator of the colon-selective Treg transcriptome (Figure 5C). To determine whether these JunB targets were expressed in a specific subpopulation of Tregs, we assessed their expression in splenic and colonic Treg subsets identified by scRNA-seq (Supplementary Figures 5F and 5G). This demonstrated that JunB-dependent genes were indeed colon-selective but were expressed at similar levels in both ROR*γ*t^+^ and ST2^+^ Tregs, indicating that the JunB plays a subset-independent role in colonic Treg programming (Figure 5D). We noted that several of the downregulated genes were involved in migration: *Ccr1* and *Ccr5* are inflammatory chemokine receptors, the latter previously shown to direct Tregs to inflamed tissues (50, 51), while *Adam8* influences cell migration through proteolytic cleavage of PSGL-1 and is also associated with invasiveness in cancer (52, 53). On the other hand, *Ccr7* was upregulated in the absence of JunB and is important for Treg migration to LN, suggesting that JunB may function to oppose lymphoid homing in Tregs (51). Thus, JunB may control migration of colonic Tregs either within or between tissues, which could partly explain defective colonic immune homeostasis in KO and iKO mice.

Aside from effects on migration-related genes, we observed that Granzyme B (*Gzmb*) was both strongly JunB-dependent and highly colon-preferential, suggesting that it might be an important target of JunB (Figures 5C and 5D). Furthermore, because Granzyme B has been described as a suppressive mechanism employed by Tregs (29–31), we reasoned that loss of Granzyme B expression could underlie unrestrained T cell proliferation in the colons of KO and iKO mice. We further assessed the likelihood that *Gzmb* was a direct target of JunB by integrating Treg chromatin accessibility [as measured by assay for transposase accessibility (ATAC)-seq] (6) and JunB chromatin immunoprecipitation (ChIP)-seq data from Th17 cells (14). This showed that JunB bound several putative regulatory elements upstream of *Gzmb*, including two that had significant ATAC signal specific for colonic Tregs – one of which was the *Gzmb* promoter (Figure 5E) – strongly implicating *Gzmb* as a direct target of JunB. We validated the JunB-dependence of Granzyme B expression using flow cytometry, which demonstrated that a small population of colonic Tregs (∼4%) expressed Granzyme B protein in iHET mice but was absent in iKO mice (Figure 5F). Unexpectedly, Granzyme B-expressing Tregs were enriched in the IL-10^+^ subset of colonic Tregs (Figure 5F), suggesting that Granzyme B is primarily expressed by Tregs actively engaging multiple effector mechanisms. Of note, scRNA-seq data indicated that *Gzmb* mRNA was expressed in ∼70% of Tregs in both ROR*γ*t^+^ and ST2^+^ subsets (Figure 5D), suggesting that additional post-transcriptional regulatory mechanisms control Granzyme B protein expression in Tregs. Together, these data demonstrate that Granzyme B is a major transcriptional target of JunB and suggest that reduced expression of Granzyme B may be responsible – at least in part – for the impaired function of colonic Tregs lacking JunB.

## Discussion

Here we defined a novel role for JunB in the control of intestinal regulatory T cell programming. JunB was essential for the maintenance of Treg-mediated immune homeostasis in the colon, which led to aberrant T helper cell cytokine responses in mice lacking JunB expression in Tregs. We identified two distinct roles for JunB in Treg biology: the maintenance of CD25^-^ Treg cells (encompassing Tfr cells) and the activation of a colonic suppression program including the expression of Granzymes A and B. The exquisite specificity of JunB for these Treg functions is underscored by the JunB-independent expression of the majority of Treg cell suppressive molecules, organ-specific immune dysfunction in mice with Treg-restricted JunB ablation, and the JunB-independence of both the general eTreg population as well as most major tissue-specific subpopulations of Treg cells. Overall, our work provides novel insight into the regulatory mechanisms underlying transcriptional programming in Tregs and shows that AP-1 TFs are an important part of the regulatory network enabling Treg effector function in non-lymphoid tissues.

The molecular drivers controlling adaptation of Treg transcriptional programming to diverse tissues have remained largely unknown. The idea that AP-1 TFs might play a tissue-specific regulatory role in Tregs was initially supported by the identification of BATF and co-regulator IRF4 as critical regulators of VAT-resident Tregs (9). However, later experiments suggested that BATF plays a more general role in the differentiation of eTregs (11), and is thus broadly required for Treg effector function independent of anatomical location. Therefore, whether canonical AP-1 TFs – such as JunB – might facilitate adaptation of Tregs to specific tissues was unclear and seemed doubtful due to the role of JunB as a major binding partner in the BATF-IRF4 complex (40–42). Our present work overturns this notion, demonstrating that JunB performs regulatory functions in Tregs that are primarily important for function in the intestine. Given that JunB exhibits variable expression in Tregs from different organs, this suggests that JunB may function to link local environmental cues to tissue-specific gene expression.

One of the critical roles we identified for JunB is the maintenance of CD25^-^ Tregs, a major Treg population present in the PP (∼50% of Foxp3^+^ cells) but also to a lesser degree in the spleen and LN. The importance of CD25^-^ Tregs is unclear: they exhibit a highly activated phenotype characterized by elevated ICOS and PD-1 expression, suggestive of chronic antigen exposure, but the majority have not been attributed any specific function – aside from the notable subpopulation that exist as Tfr cells. We confirmed that the loss of CD25^-^ Tregs following JunB deletion does not result from loss of Foxp3 expression, which is aligned with a recent report showing that CD25^-^ Tregs are not particularly unstable (34). Using RNA-seq, we found that JunB-deficiency in CD25^-^ PP Tregs leads to reduced expression of genes involved in several metabolic pathways – notably fatty acid metabolism and oxidative phosphorylation. Interestingly, Tfr cells have recently been shown to depend on mTORC1 signaling and aberrant Akt-PI3K-mTOR signaling can drive both CD25^-^ Treg accumulation and Tfr differentiation (47, 48). Furthermore, *Raptor* – a critical component of the mTORC1 complex – was shown to be important for expression of CTLA-4 and ICOS at the protein level due to its ability to drive lipid metabolism in Tregs (46). Considering that we observed reduced CTLA-4 and ICOS protein expression in the absence of JunB (despite no detectable effect on transcription of either gene), these data suggest that altered JunB-dependent metabolic pathways – in particular lipid metabolism – may be responsible for loss of CD25^-^ Tregs.

In the colon, we found that the role of JunB was to drive a subset of the colonic Treg cell transcriptome shared by both ROR*γ*t^+^ and ST2^+^ Tregs, rather than to selectively maintain either subpopulation. While we did observe a striking reduction in the frequency of colonic Tregs after JunB deletion, as well as reduced expression of several migration genes, the observation that Treg numbers in KO animals were the same or increased suggests that impaired suppression of effector T cell accumulation is the cause of reduced Treg cell frequency. Granzyme A and B are the most notable members of the JunB-dependent colonic Treg program and have been previously defined as important mediators of suppression by Treg cells (29–31, 44). scRNA-seq analysis demonstrated that Granzyme B is expressed by the majority of colonic Treg cells, including both ROR*γ*t^+^ and ST2^+^ Tregs, suggesting that Granzyme B expression represents a subset-agnostic suppressive mechanism employed by colonic Tregs. However, we found that only a small subset of colonic Tregs actively expressed Granzyme B protein at steady-state, suggesting the existence of post-transcriptional regulatory mechanisms. Nevertheless, Granzyme B^+^ Tregs were completely lost when JunB was ablated, implying that JunB is critical for cytolytic effector function in colonic Tregs. Although the roles of Granzyme A and B in maintenance of homeostasis by colonic Tregs have not been directly addressed, previous studies have demonstrated that Tregs use granzymes for suppression in models of cancer (30), respiratory infection (31), and graft-versus-host-disease (44), suggesting that this suppressive mechanism is likely to be important in the colon. Because loss of Granzyme A and B expression preceded the onset of immune dysregulation, these data suggest that loss of this suppression mechanism – potentially compounded by defects in migratory genes – likely drives the selective loss of colonic homeostasis upon JunB deletion in Tregs.

Our observations on the role of JunB in Tregs are somewhat at odds with a recent report by Koizumi *et al*. (17), although we do agree on several points. Like Koizumi, we find that ablation of JunB in Tregs using *Foxp3^YFP-cre^* results in loss of immune homeostasis and a colon-specific reduction in Treg frequencies. However, Koizumi et al. attribute this phenotype to a requirement for JunB in both eTreg differentiation and direct transcriptional activation of *Icos* and *Ctla4*, whereas our data support neither of these claims. Importantly, these mechanistic insights were surmised from experiments using *Junb^f/f^ CD4^cre^* mice and mixed bone marrow chimeras derived therefrom (17), which have since been demonstrated to be problematic models for the assessment of the role of JunB in Treg function. Specifically, *Junb^f/f^ CD4^cre^* mice have impaired thymic Treg differentiation and loss of peripheral Tregs resulting from a Treg-extrinsic defect in IL-2 production by Foxp3^-^ T cells in the thymus (14, 18). Moreover, both thymocytes and peripheral T cells from *Junb^f/f^ CD4^cre^*mice are strongly disadvantaged under competitive conditions (18) – irrespective of Foxp3 expression – highlighting an important role for JunB during T cell development or maturation that confounds the observation of JunB-deficient eTreg generation in mixed bone marrow chimeras. In contrast, our study employs multiple Treg-restricted genetic models to demonstrate that apparent JunB-dependent ICOS expression is due to loss of CD25^-^ Tregs – rather than direct regulation by JunB – and instead suggests that the importance of JunB in Tregs derives from an essential role in maintenance of both Tfr cells and a novel colonic suppression program.

In summary, our study defines JunB as a critical regulator of intestinal Treg cell functionality, with important roles distinct from those of other bZIP / AP-1 proteins, such as BATF and c-Maf. The major function of JunB in Tregs is to maintain CD25^-^ Treg cells – perhaps by facilitating metabolic adaptation – and to drive expression of a subset of genes in the colon including granzymes A and B. Remarkably, despite largely normal expression of most Treg effector genes, loss of the JunB-dependent subset of the Treg transcriptome is sufficient to cause spontaneous immune dysregulation, suggesting that impairment of organ-specific Treg effector mechanisms may drive development of autoimmune disease. Because of the wide distribution of AP-1 binding sites in putative tissue-specific regulatory elements (6), this implies that rare single-nucleotide variants (SNVs) in AP-1-responsive regulatory elements could lead to impaired TF binding that – combined with appropriate environmental triggers – may precipitate organ-specific autoimmunity by affecting context-dependent Treg effector functions. Indeed, previous studies have identified disruption of AP-1 motifs by SNVs as a potential contributor to Crohn’s disease risk (54). This is particularly important considering that most analyses of Treg cells in human autoimmunity focus on cells derived from peripheral blood, which differ substantially from their tissue-resident counterparts (55). Future efforts to evaluate the tissue-specific functions and underlying transcriptional regulatory mechanisms of human Tregs – especially those controlled by AP-1 – are therefore critical to understanding the role of Tregs in the etiology of autoimmune and inflammatory conditions.

## Acknowledgements

We thank Erwin Wagner (CNIO, Spain) for providing *Junb* conditional mice; and Alexandra Miggelbrink, Eunchong Park and Morgan Parker for technical assistance. We acknowledge the expert assistance of N. Martin and L. Martinek with flow cytometry sorting. This work was funded by a Whitehead Scholar Award (to M.C.) and by NIH grant R01 GM115474.

## Author Contributions

J.D.W. designed, performed, and analyzed experiments; performed computational analyses; and wrote the manuscript. M.C. designed experiments and edited the manuscript.

## Methods

### Mice

*Junb* floxed mice were initially obtained from Erwin Wagner (CNIO, Spain) followed by backcrossing to C57BL/6 for at least 5 generations (56). For some experiments, *Junb*-floxed mice were bred to *Foxp3^YFP-cre^* mice (Jackson Labs; Stock No. 016959) (20). Because we observed promiscuous germline deletion of *Junb* in these animals, we bred *Junb^flox/flox^* males to *Junb^flox/+^ Foxp3^YFP-cre/+^* females to minimize aberrant deletion; thus, due to X-linked inheritance of *Foxp3*, all experiments using these mice employed male *Junb^flox/+^Foxp3^YFP-cre^*(HET) or *Junb^flox/flox^Foxp3^YFP-cre^* (KO) animals, generally between 8-12 weeks of age unless otherwise specified. For experiments involving inducible deletion, *Junb*-floxed mice were bred to *Foxp3^EGFP-cre-ERT2^*mice (Jackson Labs; Stock No. 016961) (36) and also *Rosa26^STOP-ZsGreen^* (Jackson Labs; Stock No. 007906) or *Rosa26^STOP-tdTomato^*reporter mice (Jackson Labs; Stock No. 007914) to enable permanent labeling of cells that activated Cre recombinase (57). To induce deletion of *Junb*, 10-12 week old male and female mice were lightly anesthetized with Isoflurane, followed by administration of 8 mg of tamoxifen (Sigma; T5648) in 200 μL olive oil (Sigma; O1514) by oral gavage on days 0, 1, and 3. Mice that received tamoxifen were sacrificed for analysis on either day 7 or day 14. To minimize the effect of intestinal microbial variation on our results, mice in all experiments were littermates and genotypes to be compared were always co-housed.

### Isolation of cells from tissues

Spleen and lymph nodes were mechanically dissociated by grinding tissues through a 40 μm cell strainer into staining buffer (PBS supplemented with 0.5% BSA and 2 mM EDTA) or IMDM (supplemented with 10μg/mL glutamine, 10U/mL penicillin, 10μg/mL streptomycin, 10μg/mL gentamycin, and 10% FBS) if cells were to be stimulated for cytokine measurements. Red blood cells were removed from splenocytes by lysis with ACK buffer, followed by washing and resuspension in staining buffer or IMDM. Preparation of lamina propria lymphocytes from small intestine and colon was as described previously (14). Briefly, intestines were separated into small intestinal and colonic sections followed by removal of Peyer’s patches. Intestines were fileted lengthwise, cut into 2 cm pieces, and transferred into PBS containing 1 mM DTT with shaking to remove mucus. Intestine pieces then underwent two rounds of incubation in PBS supplemented with 1 mM HEPES and 5 mM EDTA at 37 °C with shaking to remove the epithelium, followed by washing with HBSS supplemented with 10% FBS. Digestion was performed by mincing tissue pieces in HBSS supplemented with 10% FBS, collagenase D (1mg/mL), DNAseI (0.1mg/mL), and dispase (0.1 U/mL) followed by incubation for 25 min at 37° C with shaking. The resulting cell suspension was filtered through a 100 μm cell strainer, followed by a discontinuous 40:80 Percoll gradient to remove large cells and debris. Cells were then collected from the interphase, washed, and resuspended in either staining buffer or IMDM as appropriate. Peyer’s patch cells were isolated by mincing excised Peyer’s patches in HBSS supplemented with 10% FBS, collagenase D (1mg/mL), and DNAseI (0.1mg/mL), followed by washing and resuspension in staining buffer or IMDM. VAT lymphocytes were isolated from epididymal adipose tissue by digestion for 1 hr with shaking at 37 °C in RPMI 1640 supplemented with 3% FBS, HEPES (1mM), collagenase D (1mg/mL), and DNAseI (0.1mg/mL), followed by centrifugation to remove floating adipocytes, red blood cell lysis using ACK buffer, and resuspension in IMDM.

### Antibodies and flow cytometry

To assess cytokine production in T cells, single-cell suspensions prepared as above were stimulated with phorbol-12-myristate-13-acetate (PMA; 50 ng/mL; Sigma) and ionomycin (500ng/mL; Sigma) in the presence of monensin (GolgiStop; BD) for 4 hours. Cells were surface stained with fixable viability dye (Thermo-Fisher) and Fc-Block (Biolegend), then fixed in 2% PFA for 15 minutes followed by permeabilization and staining with fluorochrome-conjugated antibodies in staining buffer containing 0.5% saponin. For transcription factor staining, cells were first stained with Fc-Block (Biolegend) for 10 minutes on ice, followed by fluorochrome-conjugated antibodies in staining buffer for 20 min on ice and fixation/permeablization using the Foxp3 Fix/Perm buffer set (Thermo-Fisher). Subsequent intracellular staining using fluorochrome-conjugated antibodies was performed in Permeabilization Buffer (Thermo-Fisher). Antibodies used for surface staining were as follows: CD4 (RM4-5; Thermo-Fisher), TCR*β* (H57-597; Thermo-Fisher), CD62L (MEL-14; Thermo-Fisher), CD44 (IM7; Thermo-Fisher), CD25 (PC61; Thermo-Fisher), ICOS (C398.4; Thermo-Fisher), CXCR5 (2G8; BD), PD-1 (29F.1A12; Biolegend), ST2 (RMST2-2; Thermo-Fisher), KLRG1 (2F1; Thermo-Fisher), B220 (RA3-6B2; Thermo-Fisher and BD), CD38 (90; Thermo-Fisher), anti-mouse IgG1 (for surface staining;A85-1; BD) and Fixable Viability Dye (Thermo-Fisher). Antibodies for intracellular staining were: Foxp3 (FJK-16s; Thermo-Fisher), ROR*γ*t (B2D; Thermo-Fisher), Granzyme B (GB12; Thermo-Fisher), Helios (22F6; Thermo-Fisher), CTLA-4 (UC10-4B9; Thermo-Fisher), IL-17A (eBio17B7; Thermo-Fisher), IFN*γ* (XMG1.2; Thermo-Fisher), GM-CSF (MP1-22E9; Thermo-Fisher), IL-4 (11B11; BD), IL-13 (eBio13A; Thermo-Fisher), and Bcl-6 (K112-91; BD). JunB staining was performed using unconjugated clone C-11 (Santa Cruz) followed by an anti-mouse IgG1 secondary (M1-14D12; Thermo-Fisher). Stained cells were analyzed predominantly on a BD LSRFortessa X-20 or, rarely, a BD FACSCanto II (Duke Cancer Center Flow Cytometry Core). For cell sorting, cells were stained as above but without fixation, followed by sorting on a Beckman-Coulter MoFlo Astrios at the Duke Cancer Center Flow Cytometry Core or the Duke Human Vaccine Institute Flow Cytometry Core.

### Analysis of fecal microbe-bound IgA by flow cytometry

To assess IgA binding to intestinal microbes, fresh fecal pellets were collected from mice and homogenized by vortexing with glass beads in PBS. The resulting suspension was centrifuged at low-speed (400 x *g*) to remove large particles, and the supernatant was removed to a fresh tube and centrifuged at high-speed (9000 x *g*) to pellet intestinal microbes. Microbial pellets were then resuspended in PBS containing 1% BSA, and surface stained with antibodies against mouse IgA and Ig*κ* for 20 minutes on ice. Cells were then washed with PBS and fixed in 3% formadehyde for 5 min. Fixed microbes were then resuspended in staining buffer containing DAPI, followed by acquisition on a flow cytometer.

### Library preparation and RNA sequencing

RNA was collected from sorted Tregs by lysis in Trizol, chloroform-assisted phase separation, and RNA purification from the aqueous phase using the Qiagen RNeasy-micro kit following the provided instructions. Purified RNA was submitted to the Duke Sequencing and Genomic Technologies core facility for library preparation using the Clontech SMARTer v4 Ultra-Low kit (Takara). RNA-seq libraries were sequenced on either an Illumina HiSeq4000 flow cell as 50 bp single-end reads (for *Foxp3^YFP-cre^*) or on an Illumina NovaSeq S-Prime flow cell as 50 bp paired-end reads (for *Foxp3^EGFP-cre-ERT2^*).

### RNA-seq pre-processing, read alignment, and feature assignment

Analysis of the raw NovaSeq reads using FastQC (vX; Babraham Bioinformatics) revealed a significant fraction of reverse reads containing the Clontech SMARTer adapter, thus these data were treated as single-end by omitting the reverse read in downstream analyses. Otherwise, all raw data appeared normal using standard quality metrics (FASTQC). Raw reads were trimmed to remove adapters and low-quality bases (Q < 20) using TrimGalore(v0.4.4) / CutAdapt (v1.14; Babraham Bioinformatics); reads with length < 20 bp after trimming were discarded. Alignment to the mm10 genome (GENCODE; GRCm38 primary assembly) and reference transcript annotation (GENCODE; GRCm38 vM22) was performed using STAR (v2.4.1) (58) allowing no multi-mapped reads (--outFilterMultimapNmax 1), and no novel splice junctions (--alignSJoverhangMin 500). Resulting BAM files were sorted using samtools (v1.9) (59), and reads were assigned to features in the mm10 transcript annotation using featureCounts (v1.5.3; WEHI Bioinformatics) (60) with default settings. QC metrics for all steps above were aggregated using MultiQC (v1.7) (61), which demonstrated high quality alignment and assignment, with no sample-specific biases detected. Analyses of publicly-available bulk RNA-seq data were performed from raw reads as described above. All steps of this pipeline were performed using custom scripts written for the Snakemake workflow manager (62).

### Differential expression analysis

Raw read counts were analyzed for differential expression (DE) using edgeR (v3.26.8) (63). Briefly, lowly-expressed genes were filtered out by removing those that did not have at least 1 count-per-million in at least as many samples as the size of the smallest experimental group. Counts were then normalized for library size using the trimmed mean of M-values approach implemented by edgeR. For our PP samples (both *Foxp3^YFP-cre^* and *Foxp3^EGFP-cre-ERT2^*, each having 2 samples per group) and publicly-available data, we employed a simple 2-group approach: dispersions were calculated in edgeR using estimateDisp(), followed by DE analysis using the exactTest method with prior.count set to 1. For coLP samples from *Foxp3^EGFP-cre-ERT2^*mice (3 animals per group), exploratory analysis of normalized read counts using a sample-wise Pearson correlation coefficient matrix revealed that coLP samples clustered more closely based on cage than genotype (co-housed animals clustered together), indicating that cage-specific effects influenced gene expression in coLP Tregs. However, because these data were generated in a blocked manner (one iHET and one iKO animal per cage), we included cage as a factor alongside experimental group to calculate DE using a generalized linear model (glm). Specifically, the design formula (∼group + cage) was used to calculate dispersions with estimateDisp(), which were then passed to the glmFit() function, again using a prior.count of 1. Differentially-expressed genes were then identified using the glmLRT() function in edgeR, contrasting only the iHET and iKO groups. In all comparisons, genes were considered differentially-expressed if they were attributed a Benjamini-Hochberg false discovery rate (FDR) < 0.05.

### Gene set enrichment analysis (GSEA)

GSEA (64) was performed using the Broad Institute’s Java-based GSEA application (v3.0) run in pre-ranked mode with 1000 permutations. To enrich for genes likely to be biologically-significant, DE data were first filtered to remove those genes having fewer than 5 transcripts-per-million (TPM) in at least as many samples as the size of the smallest experimental group being compared. These were then ordered by the −log10(*p*-value) multiplied by the sign of the calculated log_2_ fold-change, resulting in a ranked gene list that was used as input for GSEA. We used the curated MSigDb Hallmarks gene set collection (65) to perform exploratory analysis, with gene sets having an FDR < 0.25 and nominal *p*-values < 0.05 considered significantly enriched. All normalized enrichment scores (NES) reported were calculated using the full hallmarks collection.

### Single-cell RNA-seq (scRNA-seq) analysis

Processed scRNA-seq data (as a matrix of counts) were obtained following the link provided in the original publication (4). Seurat (v3.1.0) (66) was used for all downstream analysis. Briefly, cells were filtered to retain those having 1000 to 15000 unique transcripts corresponding to between 700 to 3500 genes per cell and to select only cells that were sorted as Tregs. Filtered data were then normalized using NormalizeData(), followed by identification of variable features [FindVariableFeatures()], and data scaling [ScaleData()] all with default settings. Scaled and normalized data were used to assess *Junb* expression in Tregs across organs without further processing (Figure 1A). For all other analyses, Treg cells obtained from spleen and colon were extracted from the full dataset, and variable feature extraction and data scaling were repeated as above. Principal component (PC) analysis was performed on the variable features using RunPCA(); 8 PC’s were chosen to be used for downstream analyses. FindNeighbors was used to compute the KNN graph using the first 8 PCs, followed by cluster identification using FindClusters() with resolution set to 0.5. Cluster markers were identified using FindAllMarkers() and were compared to known Treg subset markers to manually relabel clusters with the appropriate subset ID (Supplementary Figure 5F and 5G). Two small clusters of splenic Tregs were identified that did not appear to represent canonical Treg subsets: unique cluster markers for one of these was dominated by interferon stimulated genes (ISG), whereas cursory analysis of the other revealed no clear distinction and was thus labeled “Other”. The remaining clusters corresponded to splenic cTregs (high expression of *Sell and Ccr7*); splenic eTregs (low expression of *Sell* and *Ccr7*, elevated expression of *Icos* and *Maf*); colonic ROR*γ*t^+^ Tregs [low expression of *Ikzf2* (Helios) and *Gata3*, high expression of *Maf*, *Il10*, and *Ccr2*], colonic ST2^+^ Tregs [high expression of *Ikzf2*, *Gata3*, and *Il1rl1* (ST2)], and colonic lymphoid tissue (LT)-like Tregs (high expression of *Ccr7, S1pr1*, and *Sell*, low expression of ROR*γ*t^+^ and ST2^+^ Treg markers). *t*-SNE was performed using RunTSNE() with the first 8 PCs and visualized using DimPlot().

### ChIP-seq and ATAC-seq Analysis

Publicly-available JunB ChIP-seq (14) and colon-versus-spleen Treg ATAC-seq (6) were analyzed as follows. As with RNA-seq, raw reads were trimmed to remove adapters and low-quality bases (Q < 20) using TrimGalore (with paired-end or single-end mode as appropriate); reads with length < 20 bp after trimming were discarded. Alignment to the mm10 genome was performed using bowtie2 (v2.3.5.1) (67) in single-end mode for ChIP-seq and paired-end for ATAC-seq data. Aligned reads were sorted with samtools, filtered to remove reads overlapping genomic regions with high, anomalous signal across multiple methods (ENCODE “blacklisted” regions) (68) using bedtools (v2.29.0) (69), and duplicates were marked for exclusion in downstream analyses using Picard markDuplicates (v2.20.6; Broad Institute) (70). RPKM-normalized bigWig files for visualization were created using the bamCoverage function from deepTools (v3.3.0) (71) with the X, Y, and mitochondrial chromosomes excluded from normalization.

### Plotting and statistics

Visualizations of summary data for non-sequencing methods were prepared using the Python packages pandas, seaborn, and matplotlib. Plots related to RNA-seq were prepared in R using ggplot2, aided by ggrepel and dplyr. With the exception of DE analysis and GSEA (described above), all statistical tests were performed using the Python libraries scipy.stats and statsmodels. In general, due to the lack of normality in most data, the Mann-Whitney *U*-test was used to perform statistical comparisons. However, in certain cases where data were approximately normal and/or the number of observations were relatively high, Welch’s unequal variance *t*-test was used (with two-way ANOVA as applicable). For all methods, multiple comparisons were accounted for by adjusting *p*-values using the Holm-Bonferroni procedure.

