## Supplementary Data for "JunB controls intestinal effector programs in regulatory T cells" (from another version of this preprint: version 1, 2019-09-16)

**Supplementary Figure Legends**

**Supplementary Fig. 1. Expression of JunB in Treg subsets and correlation between JunB-dependent weight loss and proinflammatory cytokine production.**

**(A)** Expression of JunB in cTreg and eTreg cells, pre-gated as TCRβ$^{+}$CD4$^{+}$Foxp3$^{+}$ cells. Significance determined by two-way ANOVA with Tukey's HSD. **(B)** Association between colonic proinflammatory cytokine production and decreased body weight. Shown are proportions of cytokine-producing cells amongst TCRβ$^{+}$CD4$^{+}$ cells in the colon. All summary data represent mean with a bootstrapped 95% confidence interval. **p < 0.01.

**Supplementary Fig. 2. Altered Treg and cytokine responses following Treg-specific JunB deletion.**

**(A)** Total cell count from the indicated organs in HET and KO mice. Statistical significance is based on a significant ANOVA interaction term followed by planned comparisons within each organ using Welch's *t*-test with Holm-Bonferroni correction. **(B)** Frequency of KLRG1$^{+}$ cells among TCRβ$^{+}$CD4$^{+}$Foxp3$^{+}$ cells. Statistical significance determined by Mann-Whitney U-test with Holm-Bonferroni correction. **(C)** Comparison of relative CD44 expression on cTreg and eTreg cells from HET and KO mice. **(D)** Correlation of the indicated cytokines with the abundance of ST2$^{+}$ Treg cells. Spearman's ρ was calculated for pooled values from both genotypes and all organs, and indicated a significant correlation in all cases shown ($p < 1x10^{-5}$). **(E)** Frequency of Helios$^{+}$ cells among TCRβ$^{+}$CD4$^{+}$Foxp3$^{+}$ cells. Significance determined by Mann-Whitney U-test with Holm-Bonferroni correction. All summary data represent mean with a bootstrapped 95% confidence interval. *p < 0.05; **p < 0.01; ***p < 0.001; ns, not significant.

**Supplementary Fig. 3. Loss of JunB in Tregs alters germinal center responses and Treg phenotypes.**

**(A)** Frequency of PD-1$^{hi}$Bcl-6$^{+}$ or PD-1$^{hi}$CXCR5$^{+}$ Tfh cells among TCRβ$^{+}$CD4$^{+}$Foxp3$^{-}$ cells. **(B)** Frequency of CD38$^{-}$Bcl-6$^{+}$ GC B cells among B220$^{+}$ B cells. **(C)** Frequency of IgG1$^{+}$ GC B cells (defined as in *(B)*). **(D)** Frequency of CD38$^{+}$IgG1$^{+}$ memory B cells among B220$^{+}$ cells. Statistical significance determined using Mann-

Whitney U-test with Holm-Bonferroni correction. **(E)** Representative contour plots (left) and summary data (right) showing IgA and Igκ binding to DAPI+ microbes isolated from feces of Rag1-KO mice, JunB HET, and JunB KO mice. **(F)** Volcano plot of RNA-seq data comparing PP Treg cells from HET and KO mice. Genes with FDR < 0.05 are shown in blue, with select genes highlighted in orange. *p*-values were capped at $1 \times 10^{-40}$ for visualization. **(G)** Expression level of *Icos* in PP Treg cells from HET and KO mice, as determined by RNA-seq. TPM, transcripts per million. **(H)** Frequency of IL-2-expressing TCRβ+CD4+ T cells as determined by flow cytometry. Statistical significance determined by Mann-Whitney U-test with Holm-Bonferroni correction. **(I)** Representative contour plot showing the relationship between CD25 and ICOS expression. **(J)** Relative expression of ICOS on CD25+ and CD25- Treg cell subsets. Significance determined by Mann-Whitney U-test with Holm-Bonferroni correction. All summary data represent mean with a bootstrapped 95% confidence interval. *p < 0.05; **p < 0.01.

**Supplementary Fig. 4. Inducible deletion of *Junb* recapitulates key phenotypes of JunB-KO Tregs.**

**(A)** Activity of *Foxp3*$^{YFP\text{-}cre}$ across hematopoietic lineages, assessed by FACS as the proportion of ZsGreen+ cells within the indicated cell type. Data shown are from one experiment. **(B)** Efficiency of tamoxifen-induced Cre activity, assessed by flow cytometry as the proportion of Foxp3+ cells that were labeled with ZsGreen 14 days after tamoxifen administration. **(C)** Number of Foxp3+ZsGreen+ cells across organs in iHET and iKO mice. **(D)** Total cell number in the indicated organs of iHET and iKO mice. **(E)** Frequency of eTreg cells among Foxp3+ZsGreen+ cells (left) and relative gMFI of CD44 expression on eTregs (right). Statistical signifiance determined by Welch's *t*-test. **(F)** Frequency of RORγt+ cells among Foxp3+ZsGreen+ cells determined by flow cytometry. Statistical significance determined by Mann-Whitney U-test with Holm-Bonferroni correction. **(G)** Frequency of ST2+ cells among Foxp3+ZsGreen+ cells determined by flow cytometry. **(H)** Frequency of IL-2-expressing cells among TCRβ+CD4+ZsGreen- cells. Statistical significance determined by Mann-Whitney U-test with Holm-Bonferroni correction. **(I)** Relative gMFI of ICOS on Foxp3+ZsGreen+ cells. Statistical significance determined by Mann-Whitney U-test with Holm-Bonferroni correction. **(J)** Relative gMFI of PD-1 on Foxp3+ZsGreen+ cells. Statistical significance determined by Mann-Whitney U-test. **(K)** Transcripts per million (TPM) of select effector genes in PP Tregs as determined by RNA-seq, orange asterisks indicate genes identified as significantly different by DE analysis (FDR < 0.05). All summary data represent mean with a bootstrapped 95% confidence interval. *p < 0.05; **p < 0.01; ***p < 0.001; ns, not significant.

**Supplementary Fig. 5. Regulation of gene expression in $CD25^-$ PP Tregs and colonic Tregs by JunB.**

**(A)** Frequency of Tregs determined by FACS at day 7 post-tamoxifen administration. **(B)** Expression of JunB in $TCR\beta^+CD4^+Foxp3^+ZsGreen^+$ Treg cells at day 7 post-tamoxifen, determined by flow cytometry. **(C)** GSEA analysis of RNA-seq data comparing $CD25^-$ PP Tregs from iHET and iKO mice. Pathways in blue are underrepresented in iKO, whereas those in orange are overrepresented. All pathways that reached statistical significance are shown (FDR < 0.25, *p*-value < 0.05). **(D)** GSEA analysis of RNA-seq data comparing Tfr cells and Treg cells. Pathways in orange are associated with Tfr cells, whereas those in blue are associated with non-Tfr Tregs. All pathways that reached statistical significance are shown (FDR < 0.25, *p*-value < 0.05). **(E)** Volcano plot showing differential expression in colonic Tregs from iHET vs iKO mice at day 7 post-tamoxifen. Genes with significant DE (FDR < 0.05) are highlighted in orange, with select genes labeled. **(F)** *t*-SNE plot showing clusters of cells from scRNA-seq used to assess the distribution of JunB-dependent gene expression across Treg subsets. [Interferon-stimulated genes (ISG); lymphoid-tissue (LT)] **(G)** Dot plot showing the expression of canonical markers used to assign cell identities to clusters from scRNA-seq as in *(F)*. All summary data represent mean with a bootstrapped 95% confidence interval.

### Supplementary Figure 1

A

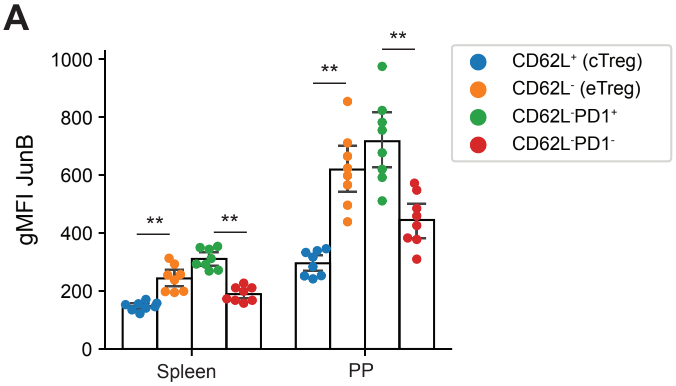


B

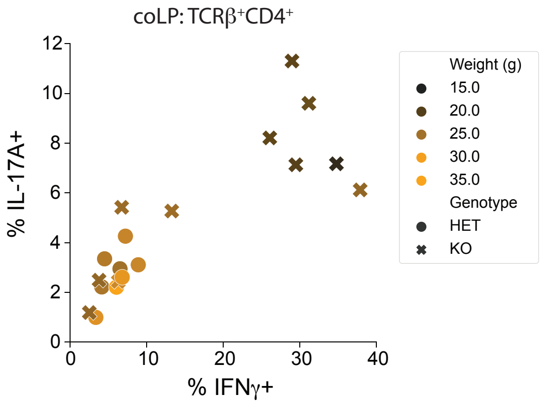

### Supplementary Figure 2

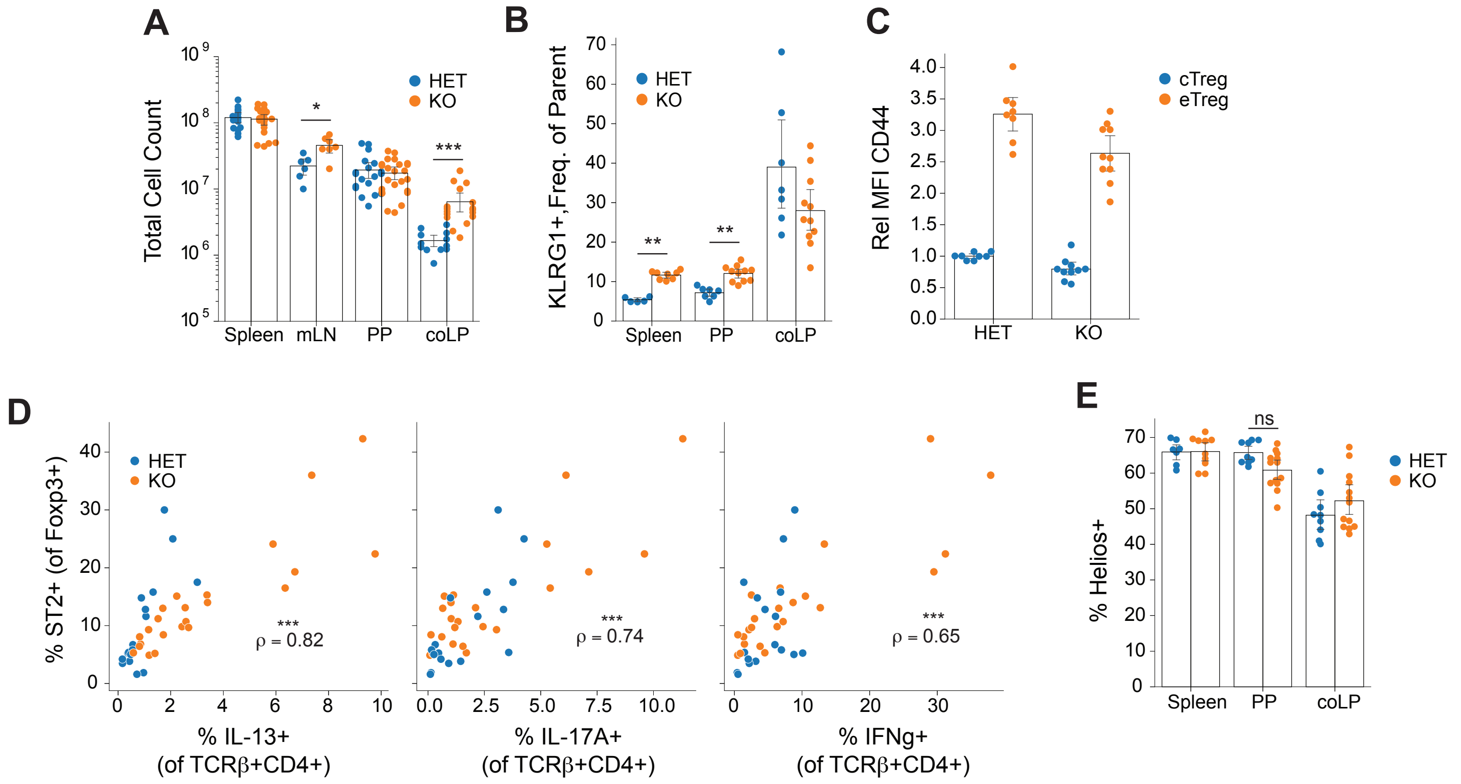

### Supplementary Figure 3

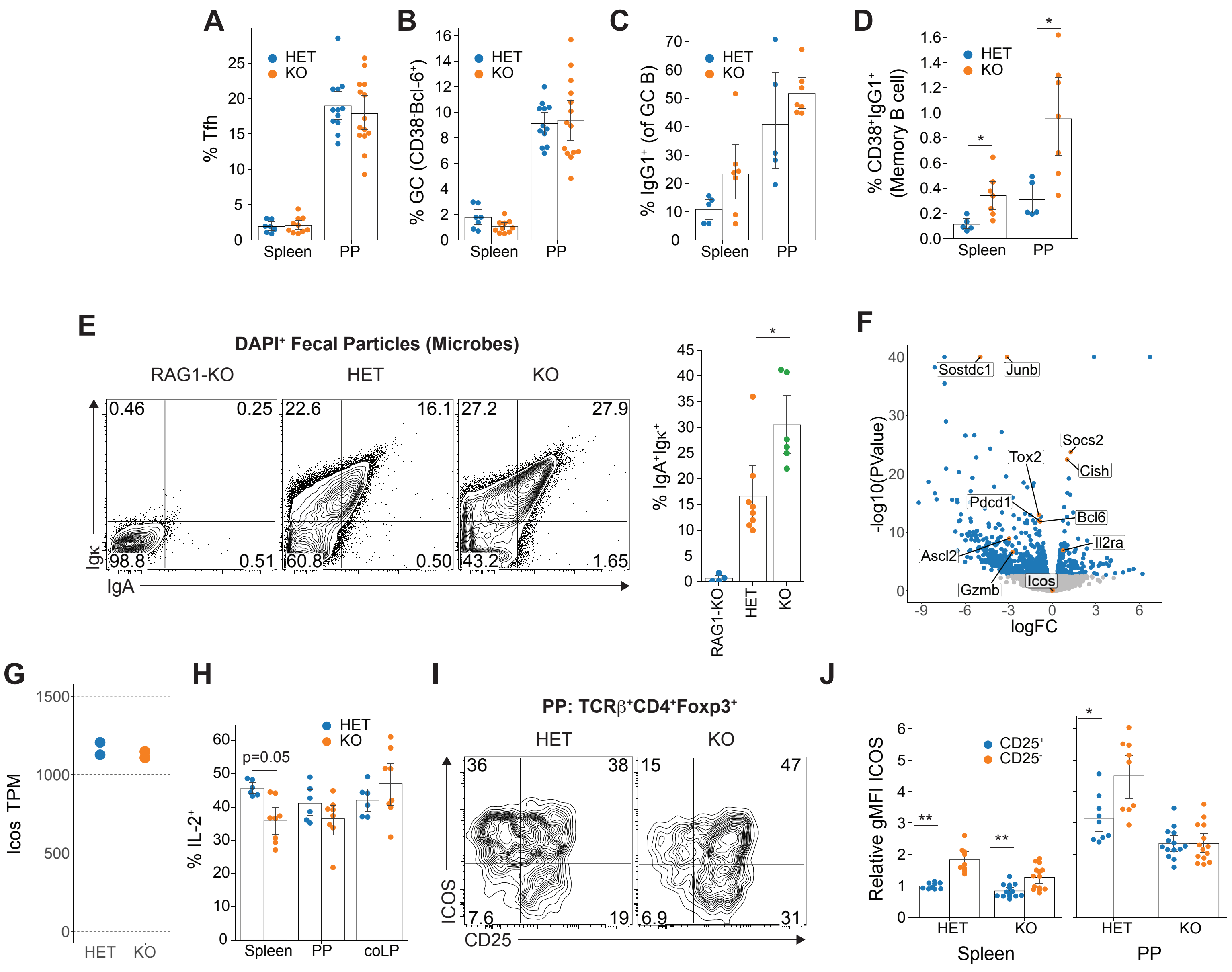

### Supplementary Figure 4

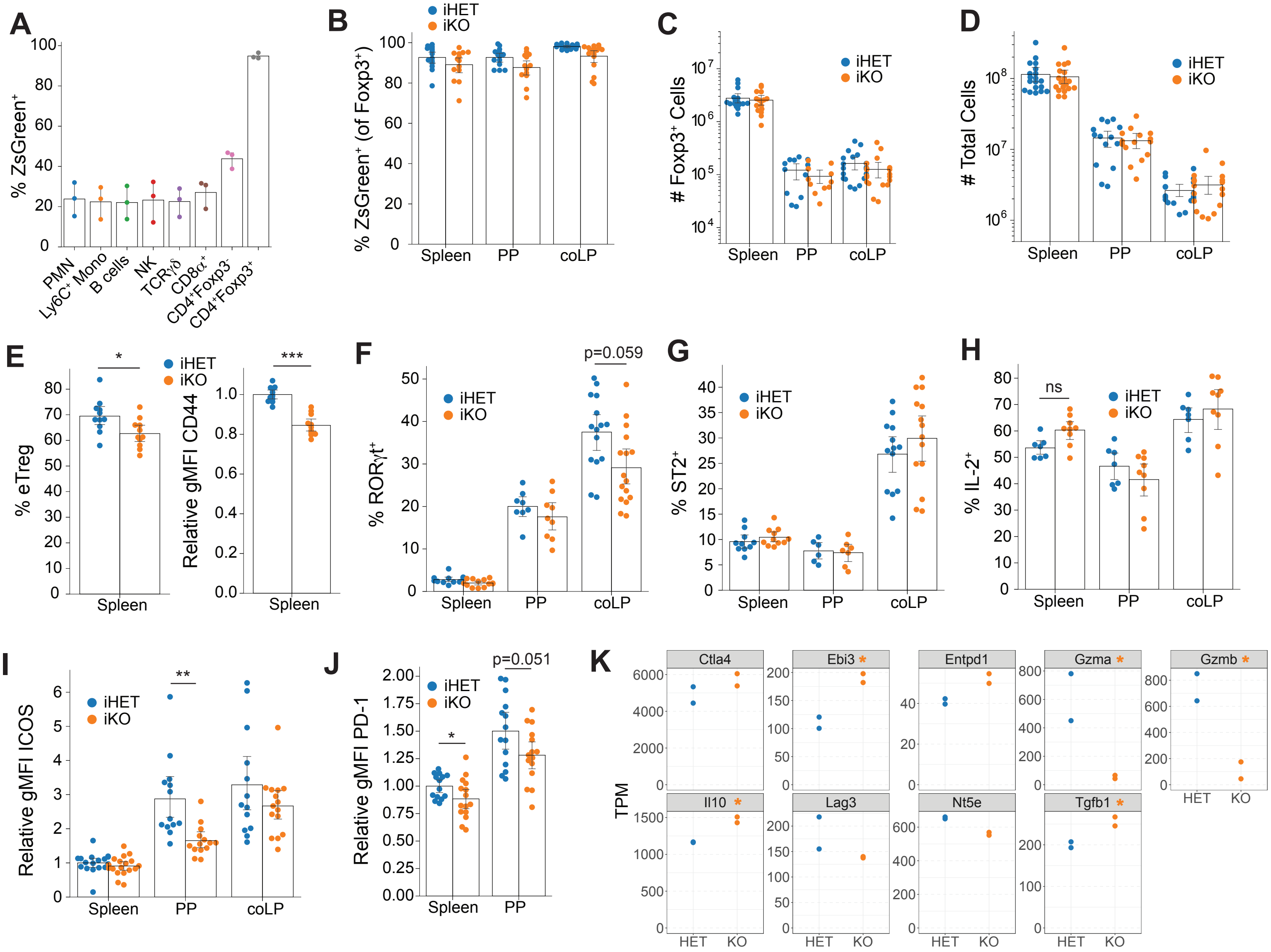

### Supplementary Figure 5

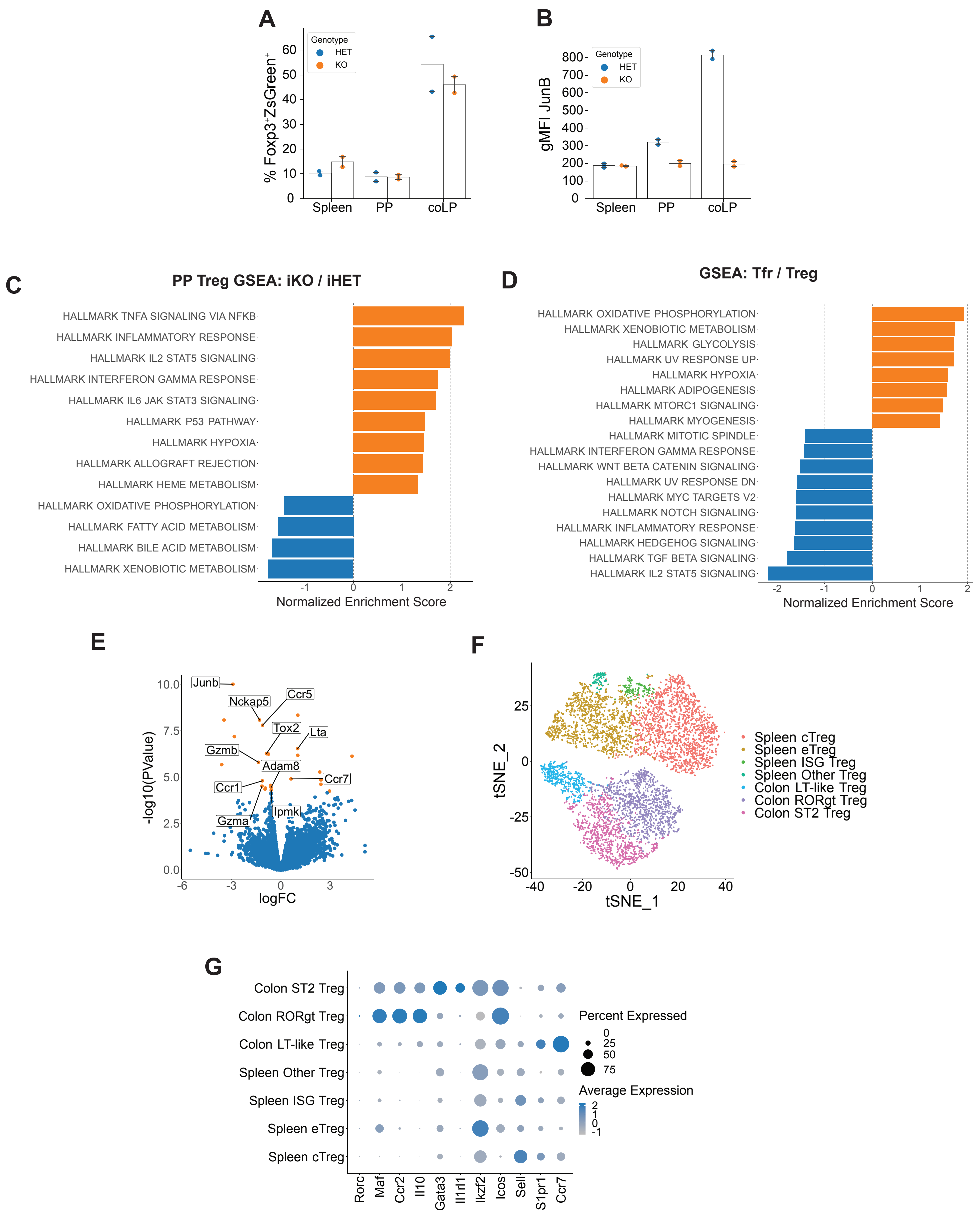
